# Exome sequencing of bulked segregants identified a novel *TaMKK3-A* allele linked to the wheat *ERA8* ABA-hypersensitive germination phenotype

**DOI:** 10.1101/784652

**Authors:** Shantel A. Martinez, Oluwayesi Shorinola, Samantha Conselman, Deven See, Daniel Z. Skinner, Cristobal Uauy, Camille M. Steber

**Author notes:** Co-first authors contributed equally to this work (Shantel A. Martinez, Oluwayesi Shorinola). Co-last authors contributed equally to this work (Cristobal Uauy, Camille M. Steber). Correspondence: Camille M. Steber e. **Author contribution statement:** SAM, OS, CMS, and CU designed the experiments. SAM and CMS developed the populations. SAM and SC performed the Louise/ZakERA8 QTL analysis. DZS and DS conducted GBS and analyzed the raw sequence data. SAM performed germination assays for the bulked segregant analysis, fine mapped *ERA8*, and introgressed *ERA8* into breeding lines. OS performed the exome capture, BSA, and designed KASP markers. SAM and OS conducted the RNA sequencing and *TaMKK3-A* alignment. CMS and CU provided resources. SAM and CMS wrote the manuscript. All authors read and approved the final submission.

## Abstract

Preharvest sprouting (PHS) is the germination of mature grain on the mother plant when it rains before harvest. The *ENHANCED RESPONSE TO ABA8* (*ERA8*) mutant increases seed dormancy and, consequently, PHS tolerance in soft white wheat ‘Zak’. *ERA8* was mapped to chromosome 4A in a Zak/‘Zak*ERA8’* backcross population using bulked segregant analysis of exome sequenced DNA (BSA-exome-seq). *ERA8* was fine-mapped relative to mutagen-induced SNPs to a 4.6 Mb region containing 70 genes. In the backcross population, the *ERA8* ABA hypersensitive phenotype was strongly linked to a missense mutation *TaMKK3-A-G1093A* (LOD 16.5), a gene associated with natural PHS tolerance in barley and wheat. The map position of *ERA8* was confirmed in an ‘Otis’/Zak*ERA8* but not in a ‘Louise’/Zak*ERA8* mapping population. This is likely because Otis carries the same natural PHS susceptible *MKK3-A-A660^S^* allele as Zak, whereas Louise carries the PHS tolerant *MKK3-A*-C660^R^ allele. Thus, the variation for grain dormancy and PHS tolerance in the Louise/Zak*ERA8* population likely resulted from segregation of other loci rather than segregation for PHS tolerance at the *MKK3* locus. This inadvertent complementation test suggests that the *MKK3-A-G1093A* mutation causes the *ERA8* phenotype. Moreover, *MKK3* was a known ABA signaling gene in the 70-gene 4.6 Mb *ERA8* interval. None of these 70 genes showed the differential regulation in wild-type Zak versus *ERA8* expected of a promoter mutation. Thus, the working model is that the *ERA8* phenotype results from the *MKK3-A-G1093A* mutation.

**Key Message:** Using bulked segregant analysis of exome sequence, we fine-mapped the ABA hypersensitive mutant *ERA8* in a wheat backcross population to the *TaMKK3-A* locus of chromosome 4A.

## Introduction

Preharvest sprouting (PHS) is the germination of mature grain on the mother plant before harvest under rainy or humid conditions (reviewed by Rodríguez et al. 2015). Lack of grain dormancy accounts for 60 - 80 % of the variation for preharvest sprouting susceptibility in bread wheat (*Triticum aestivum* L.) (DePauw and McCaig 1991). Grain dormancy refers to the inability to germinate even under favorable environmental conditions (reviewed in Bewley et al. 2013). Grain is most dormant at physiological maturity and then loses dormancy through a period of dry storage called after-ripening, through imbibition in the cold (called cold stratification), or through scarification of the seed coat (Paterson et al. 1989; Gerjets et al. 2010). Seed coat-imposed dormancy is relieved by scarification of the seed coat, whereas embryo dormancy is not. Wheat can have both forms of seed dormancy (Schramm et al. 2010).

A seed is considered germinated when the radicle or any other part of the embryo emerges from the seed coat (reviewed in Bewley et al. 2013; Mares and Mrva 2014). Hydrolytic enzymes like alpha-amylase are induced during germination to mobilize stored reserves for use in seedling growth. Alpha-amylase degrades starch during germination. Even before grain is visibly sprouted, alpha-amylase activity in dough can result in poor product end-use quality. While higher seed dormancy prevents PHS, too much dormancy at planting causes problems with uneven or poor seedling emergence (reviewed by Rodríguez et al. 2015). Thus, it is important to understand the genetic control of wheat seed dormancy and germination to develop varieties with an appropriate balance between dormancy and emergence.

Multiple environmental and genetic factors control the degree of seed dormancy. In wheat, seed dormancy is higher if the environment is cool and dry during embryo maturation, and lower if conditions are hot during embryo maturation (Nakamura et al. 2011; Mares and Mrva 2014). Plant hormone signaling controls seed dormancy and germination (reviewed in Finkelstein et al. 2008). The hormone abscisic acid (ABA) induces seed dormancy during embryo maturation whereas gibberellin A (GA) stimulates seed germination. ABA establishes seed dormancy during embryo maturation, maintains dormancy in mature seeds, and inhibits the germination of mature seeds when exogenously applied. Thus increased ABA signaling or decreased GA signaling is associated with higher seed dormancy. Dormancy is also higher in wheat varieties with red grain coat color than white (Himi et al. 2002, 2011). ABA and grain coat color can act independently and in concert to increase wheat grain dormancy (Himi et al. 2002). Mutant studies have shown that loss of ABA sensitivity can decrease seed dormancy in red wheat and that increased ABA sensitivity can increase seed dormancy in both red and white wheat varieties (Flintham 2000; Torada and Amano 2002; Schramm et al. 2010, 2012, 2013). Since white wheat varieties have lower seed dormancy and PHS tolerance, we have identified ethyl methanesulfonate (EMS)-induced genetic variants with increased ABA sensitivity as a method to increase grain dormancy and PHS tolerance in white wheat.

The EMS-induced semi-dominant *ERA8* (Enhanced Response to ABA8) mutation was isolated in the soft white spring wheat cultivar ‘Zak’ (Kidwell et al. 2002; Schramm et al. 2013). The *ERA* mutants were identified by screening for increased sensitivity to inhibition of seed germination by ABA. The ‘Zak*ERA8’* line has strong ABA sensitivity and seed dormancy that is lost rapidly through after-ripening (Schramm et al. 2013; Martinez et al. 2014, 2016). Moreover, Zak*ERA8* had higher PHS tolerance in spike-wetting tests (Martinez et al. 2016). Thus, mapping of the *ERA8* mutation would make it easier to deploy as a source of PHS tolerance in white or red wheat. ABA signaling can be controlled both at the level of hormone accumulation and at the level of the ABA sensitivity. Endogenous embryo ABA measurements showed that ABA content decreased similarly with after-ripening of *ERA8* and wild-type (WT) Zak, suggesting that the phenotype resulted from increased ABA signaling rather than decreased ABA turnover (Martinez et al., 2016). Thus, we hypothesize that *ERA8* is defined by a gain-of-function mutation in a positive regulator of ABA signaling.

Fine mapping and map-based cloning of wheat genes is challenging due to the very large size (∼17 Gb) and high redundancy of the allohexaploid bread wheat genome (Yan et al. 2003; Gupta et al. 2008; Shewry 2009). The large size is due to the fact that it contains over 80 % noncoding DNA, and that it is derived from three diploid progenitors that each contributed seven homoeologous chromosomes referred to as the A, B, and D genomes (2n = 6x = 42, genomes AABBDD).

Quantitative trait loci (QTL) for PHS and dormancy have been identified throughout the allohexaploid wheat genome (Zanetti et al. 2000; Ogbonnaya et al. 2008; Munkvold et al. 2009; Kulwal et al. 2012; Jaiswal et al. 2012; Zhou et al. 2017; Martinez et al. 2018). While homologues of ABA signaling genes have been found on wheat chromosomes 3A, 4A, and 5A, only *Viviparous-1*/*ABA-insensitive3* (*VP1/ABI3*) has been linked to differences in wheat PHS tolerance (Nakamura and Toyama 2001; McKibbin et al. 2002; Nakamura et al. 2007). To date, 18 dormancy and 13 PHS studies have identified a QTL on the long arm of chromosome 4A called the *Phs1* locus (Liu et al. 2008, 2014; Mares and Mrva 2014; Shorinola et al. 2016; Martinez et al. 2018; Zuo et al. 2019). This 4A QTL has been cloned as the wheat *Mitogen-Activated protein Kinase Kinase 3* gene (*TaMKK3-A*) (Nakamura et al. 2016; Torada et al. 2016; Shorinola et al. 2017). Another major PHS tolerance QTL on chromosome 3A was cloned from Japanese and U.S. germplasm, the wheat gene *MOTHER OF FLOWERING TIME AND TFL1* (*TaMFT1*) (Nakamura et al. 2011; Liu et al. 2013b).

The reference genome sequence of allohexaploid wheat landrace Chinese Spring has enabled the application of high-throughput sequencing methods to mapping and cloning strategies in polyploid wheat (IWGSC et al. 2018; http://www.wheatgenome.org). Even so, the large size and high redundancy of the wheat genome is still a challenge. Several methods have been employed to reduce the wheat’s genome complexity by targeted enrichment and sequencing of relevant regions of the wheat genome. Genotyping-by-sequencing (GBS) entails restriction enzyme digests of genomic DNA to enrich for DNA fragments that begin with common restriction sites for high-throughput genome resequencing (Poland et al. 2012; Truong et al. 2012). Exome sequencing (exome-seq), on the other hand, enriches specifically for the gene-containing regions of the genome through affinity capture and purification of the predicted wheat coding regions for high throughput genome resequencing (Allen et al. 2013; Henry et al. 2014; Uauy et al. 2017; Krasileva et al. 2017; Mo et al. 2017).

Bulked segregant analysis (BSA) of genome resequencing data is a powerful approach for mapping and cloning. In this approach, pooled-DNA samples from plants with contrasting phenotypes are sequenced to identify polymorphisms co-segregating with the trait of interest (Michelmore et al. 1991). In organisms with smaller genomes, it is now common to clone mutagen-induced polymorphisms using BSA by contrasting whole genome DNA sequence of bulked wild-type and mutant backcross populations (reviewed in Thole and Strader 2015). However, the wheat genome is too large for this approach. Wheat genes have been fine mapped by BSA using RNA-seq instead of DNA-seq to enrich for the expressed portion of the genome (Trick et al. 2012; Liu et al. 2012; Ramirez-Gonzalez et al. 2015a). The RNA-seq approach has the drawback that relative gene expression levels vary in different tissues and times in development, leading to the possibility that the gene of interest might be underrepresented in the sampled tissue. Exome capture has enabled affinity-purification of the predicted wheat coding regions before performing whole genome DNA resequencing (Allen et al. 2013; Henry et al. 2014; Uauy et al. 2017; Krasileva et al. 2017; Mo et al. 2017). This approach has been used to characterize the genome-sequence of EMS-mutagenized wheat TILLING populations (Harrington et al. 2019). Since this method was effective in identifying EMS-induced mutations responsible for a phenotype, this study used exome capture in a BSA-exome sequencing approach to map EMS-induced mutations linked to the *ERA8* allele.

In this study, *ERA8* was localized to chromosome 4A in a backcross population using bulked segregant analysis to map relative to EMS-generated polymorphisms characterized by high throughput DNA sequencing of exome genomic DNA (BSA-exome-seq). Fine-mapping with a larger backcross population was used to co-localize *ERA8* with mutations in and near the *TaMKK3-A* gene that was previously cloned as a gene regulating wheat preharvest sprouting tolerance. QTL analysis of RIL populations was used to confirm the gene location. This information is being used to introgress *ERA8* into soft white winter wheat to improve PHS tolerance by increasing ABA sensitivity.

## Materials and methods

### Plant material

Allohexaploid white spring wheat cultivars (*Triticum aestivum* L.) were used in this study. ‘Zak*ERA8*’ (PI 669443) is derived from the soft white spring cultivar ‘Zak’ (PI 612956) via an EMS-induced mutation resulting in enhanced response to ABA during seed germination (Kidwell et al. 2002; Schramm et al. 2013; Martinez et al. 2014). Soft white spring ‘Louise’ (PI 634865) was selected from the cross ‘Wakanz’/ ‘Wawawai’ (Kidwell et al. 2006b). Hard white ‘Otis’ (PI 634866) is a selection from the cross ‘Idaho 377s’ /3/‘Tanager S’ /‘Torim 73’//‘Spillman’(Kidwell et al. 2006a). Mapping populations derived from these lines were grown in a controlled greenhouse environment with supplemental lighting to obtain a 16 h day/8 hr night photoperiod with 15-17 °C night and 21-23 °C day temperatures.

Backcross (BC) populations were developed by three successive crosses of the ABA hypersensitive Zak*ERA8* to the mutagenesis parent Zak (Fig. S1; Schramm et al. 2013). After the third consecutive cross, a single BC_3_F_1_ line was advanced to BC_3_F_2_ by self pollination. Because *ERA8* is semi-dominant, homozygous *ERA8*/*ERA8* and +/+ BC_3_F_2_ were identified by progeny-testing in the F_3_ generation based on a non-segregating ABA-hypersensitive *ERA8*-like and normal Zak-like germination phenotype. *ERA8* and Zak genomic DNA was prepared from leaf tissue of the BC_3_F_3_ generation, and used for bulked segregant analysis. Additional BC_3_F_2:3_ seeds (X5.1 and X5.2) were phenotyped and used for fine mapping.

Two populations were constructed to confirm the *ERA8* map position, a Louise/Zak*ERA8* F_2_:F_5_ recombinant inbred line (RIL) population and an Otis/Zak*ERA8* F_2_:F_3_ population. The Louise/Zak*ERA8* RIL population was derived from F_2_ seeds from ten independent F_1_ plants from the cross of Louise to Zak*ERA8* (Louise as the female parent). The population was developed without selection through single seed descent of 225 RILs from the F_2_ generation to the F_5_ generation. Genomic DNA was extracted from single F_5_ generation plants grown in 3 L pots in the Plant Growth Facility at Washington State University, Pullman, WA. F_6_ seeds were harvested at physiological maturity (environment 1 or E1). For each RIL, 30 F_6_ seeds from E1 were advanced in headrows at the Washington State University Spillman Research Farm, Pullman, WA in 2014 and 2015 to produce F_7_ grain from the E2 and E3 environments, respectively. To construct an Otis/Zak*ERA8* F_2_:F_3_ mapping population, Otis was pollinated with Zak*ERA8* (Kidwell et al. 2006a). F_1_ seeds were advanced in the greenhouse and F_2_ harvested at physiological maturity. Genotyping was performed using genomic DNA prepared from F_2_ leaf tissue, whereas F_3_ seeds were phenotyped by ABA germination assay. Genomic DNA was extracted from Louise/Zak*ERA8* RILs and Otis/Zak*ERA8* F_2_ plants on the Qiagen BioSprint-96. DNA concentrations were determined using the BioTek Gen5 plate reader, and DNA was diluted to 50 ng/µL.

### Seed germination assays

Plating assays were used to compare ABA sensitivity during seed germination using experimentally determined conditions that best differentiated between wild type (WT) and *ERA8*/*ERA8* parents for each population or after-ripening time point (Fig. S2). The wheat grain is a caryopsis because it includes a pericarp, but is referred to here as a “grain” or “seed” for the sake of simplicity (reviewed in Bewley et al. 2013). Seeds were harvested at physiological maturity, hand threshed, then dry after-ripened at ambient temperature (20-23 °C) and humidity (15-30 %) for 5-7 weeks (as in Martinez et al. 2016). Seeds (30-90) were sterilized in a 50 mL tube with 20 mL of 10 % bleach/ 0.01 % SDS for 15 min, then rinsed 3x with 40 mL of sterile deionized water. Thirty seeds were plated on Petri dishes containing a 9-cm blue germination disk (Anchor Paper Co) saturated with 6 mL of 5 mM MES buffer, pH 5.5 (2-(N-morpholino)ethanesulfonic acid, Sigma-Aldrich) containing 2 µM, 5 µM, or 10 µM (+/-)-ABA (PhytoTechnology Laboratories). Stock solutions of 0.1 mM (+/-)-ABA were prepared in ethanol, and diluted as indicated.

Population parents were plated on varying ABA concentrations weekly from 3-8 weeks of after-ripening to determine the after-ripening time point and ABA concentration with the optimal difference between parents (Fig. S2). Seeds were stored at −20 °C for no more than 4 months to preserve dormancy until the optimal dry after-ripening time point was selected (Paterson et al. 1989). However, the larger X5 BC_3_F_3_ populations used for fine mapping were stored for 3 to 17 months at −20 °C, then the after-ripened seeds were plated on the optimal ABA concentration as indicated (Table S1). Because different wheat seed lots have different degrees of initial dormancy, and because grain after-ripens slowly in the freezer, different ABA concentrations were needed to score the phenotype in seed lots stored for different lengths of time at −20°C (Paterson et al. 1989; Schramm et al. 2013). Germination was scored daily over 5 days of incubation at 30°C. Seeds were considered germinated upon embryonic root emergence. Germination index (GI), a value weighted for speed of germination, was calculated over 5 days of imbibition as (5 x g_day1_ + 4 x g_day2_ … + 1 x g_day5_)/(5 x n) where g is the number of additional seeds germinated on each day and n is the total number of seeds (Walker-Simmons 1987; Schramm et al. 2013). GI ranges from 0-1, where a GI of 1 indicates that all of the seeds germinated on day 1 of imbibition. Raw means and standard deviations were calculated from three technical replicates of ten seeds. Chi-squared analysis was used to determine goodness-of-fit to Mendelian segregation models for F_2_:F_3_ seeds as in Schramm et al. (2013; Table S2). The χ^2^ statistic was calculated as LJ [(O – E)^2^ / (E)] where O was the observed number of seeds germinated or ungerminated, and E is the expected number of seeds germinated or ungerminated based on Mendelian segregation. A χ^2^ distribution table was used to determine the p-values based on the χ^2^ statistic and the degrees of freedom = 1. The model fits the observed values when p > 0.05, and does not fit when p < 0.05.

**Table 1.** Significant QTL found in the Louise/Zak*ERA8* RIL population.

| QTL Name | Marker | Chrm | Pos<br>(cM) | Start<sup>a</sup> | End<sup>a</sup> | SNP<br>Position | LOD<sup>b</sup> | Trait | Favorable<br>Allele<sup>c</sup> |
| --- | --- | --- | --- | --- | --- | --- | --- | --- | --- |
| *QABA.wsu-2D* | A180 | 2D | 198.9 | 36,271,198 | 36,271,099 | 54 | 5.69** | E2 D3 | <u>C</u>/T |
| *QABA.wsu-4A.1* | A1512 | 4A | 194.9 | 43,348,134 | 43,348,233 | 60 | 4.96* | E3 D1 | G/<u>A</u> |
|  | A11335 | 4A | 199.7 | 83,469,184 | 83,469,281 | 85 | 5.32* | E3 D1 | C/<u>T</u> |
| *QABA.wsu-4A.2* | A16612 | 4A | 224.2 | 238,226,439 | 238,226,538 | 35 | 5.05* | E2 D1 | G/<u>C</u> |
| *QABA.wsu-4A.3* | A28975 | 4A | 266.8 | 29,794,783 | 29,794,714 | 27 | 4.79** | E1 D2 | <u>C</u>/T |
|  | A29203 | 4A | 269.4 | 29,794,783 | 29,794,684 | 27, 88 | 4.69* | E1 D2 | <u>C</u>/T |
| *QABA.wsu-4B* | A2984 | 4B | 470.3 | 673,071,467 | 673,071,398 | 11, 15 | 4.16* | E2 D2 | C/<u>T</u> |
| *QABA.wsu-7B.1* | A8640 | 7B | 285.8 | 156,437,282 | 156,437,351 | 43 | 4.79* | E2 D1 | C/<u>T</u> |
| *QABA.wsu-7B.2* | A29262 | 7B | 365.7 | 41,637,734 | 41,637,803 | 11 | 4.71* | E2 D1 | C/<u>T</u> |

<sup>a</sup> The GBS markers was aligned to the RefSeqv1.0 reference genome and the start and end nucleotide position is reported with the SNP position within the sequence.
<sup>b</sup> Significant LOD scores with a *p-value* < 0.05 and *p-value* < 0.10 are indicate with \*\* and \*, respectively
<sup>c</sup> Favorable alleles (underlined) decrease germination percent or index. Louise or *ERA8* contributed the first or second allele, respectively.

### Exome capture and sequencing

High quality genomic DNA from the Zak/Zak*ERA8* BC_3_F_3_ bulked populations was extracted using a phenol/chloroform-based DNA extraction method followed by RNAse A digestion (as in Yu et al. 2017). A 2.5 cm long leaf sample was collected from BC_3_F_3_ plants and ground together with lines that comprised each bulk. For each individual parent, a 10-15 cm long leaf sample was ground for DNA extraction.

Whole genome DNA resequencing of the large wheat genome (16 Mb) for bulked segregant analysis is prohibitively costly (Uauy et al. 2017). Exome capture was used to restrict genomic DNA sequencing to those regions encoding predicted genes (Krasileva et al. 2017). Library preparation and sequencing was carried out at the Earlham Institute, UK as described by Harrington et al. (2019). Separate barcoded libraries were prepared from the *ERA8* parent (*ERA8/ERA8*), wild-type Zak parent (+/+), the bulked *ERA8*-like, and bulked Zak-like DNA with average insert sizes of 327, 345, 348, and 340 bp, respectively. The four libraries were multiplexed and captured on one exome-capture probeset using the wheat NimbleGen target capture following the SeqCapEZ protocol v5.0 (Krasileva et al 2017). Sequencing was conducted on a single Illumina HiSeq 2500 lane producing 125-bp paired-end reads.

Read quality was analyzed using FastQC and the reads with a base quality score ranging from 20-40 were retained (Table S3; Babraham Bioinformatics 2012). Adapter sequences were removed using Trimmomatic with the following parameters: ILLUMINACLIP (Truseq3-PE), CROP (125 bp), and MINIMUMLENGTH (50 bp) options (Bolger et al. 2014). The short read alignment tool, Bowtie2, was used to align trimmed reads to the wheat genome assembly, IWGSC RefSeq v1.0 from Chinese Spring, with some modifications (Langmead et al. 2009; Li and Durbin 2009; IWGSC et al. 2018). First, we removed the unassigned chromosome (ChrUn) containing unanchored scaffolds in this assembly as these scaffolds lack positional information useful for mapping. Also, *TaMKK3-A*, an important gene on wheat chromosome 4A that controls PHS and dormancy in wheat is unanchored in the RefSeq v1.0 assembly (IWGSC et al. 2018), so we complemented the modified RefSeq v1.0 assembly with a 30.4 kb scaffold from the Synthetic W7984 wheat genome assembly (Chapman et al. 2015) spanning the *TaMKK3-A* region. The Bowtie2 end-to-end alignment algorithm was used with the sensitivity of the alignments increased by adjusting the minimum score threshold parameter (-score-min) to L, 0, 0.15 to allow only 2-3 mismatches per alignment depending on the base quality score. The raw sequence alignment map (SAM) files were processed (converted, sorted and indexed) using SAMtools (Li et al. 2009a). To improve the accuracy of read depth values, duplicate reads resulting from PCR amplification during library preparation were removed using the Picard Markduplicate tool (http://broadinstitute.github.io/picard). Only properly paired reads, defined as correctly oriented paired reads with mapping distance less than average insert size, were retained.

### Bulked segregant analysis

Exome variants including single nucleotide polymorphism (SNP) and insertion or deletion (InDel) sites were first identified between the Zak WT and *ERA8* parents using a Samtools (mpileup)-bcftools (view) variant-calling pipeline (Li et al. 2009a). Snpsift (snpEFF) was used for the analysis, manipulation, and interactive filtering of the VCF files (Cingolani et al. 2012). First, SNP variant sites with the non-reference allele in Zak were excluded as these mostly likely represent intervarietal SNPs between Chinese Spring and Zak. Second, *ERA8*-specific SNP sites with alternate (non-reference) alleles in *ERA8* and reference allele in Zak WT were selected. Only SNPs with a variant quality score (“QUAL”) above 20 were selected to improve accuracy. Subsequent analysis concentrated on the 1,854 transition variants (C to T or G to A) as these were likely caused by EMS mutagenesis. Variants in the 1,854 sites between the bulked *ERA8*-like and WT-like sequence were called using the same variant-calling pipeline but with an additional flag to specify the position of the target sites in the mpileup analysis (--position). Bulked segregant analysis was performed to identify sites enriched in the *ERA8*-like versus WT-like bulked DNA. To do this, the *ERA8*-specific alternate allele frequencies at the selected sites were first calculated using the bulked allele frequency formula, BF = DV/DP, where DV is the number of reads with alternate allele and DP is the total read depth. To improve accuracy, analysis was restricted to sites with DP >=5. The bulked allele frequency difference (BFD) was then calculated between the two bulks as BFD = BF_ERA8_b_ - BF_ZWT_b_. The analysis was based on the rationale that *ERA8* plants should show an enriched frequency of the *ERA8*-like sequence variants in regions linked to the *ERA8* gene, but should show random segregation of unlinked sequence variants. The BF and BFD across the whole genome was visualised with Circos to highlight the *ERA8*-linked region with the highest BFD in the genome (Krzywinski et al. 2009).

### Fine mapping

*ERA8* was fine-mapped relative to EMS-induced SNPs genotyped using Kompetitive Allele Specific PCR (KASP) in Zak/Zak*ERA8* BC_3_F_2:3_ lines (Smith and Maughan 2015). KASP markers for SNPs identified through whole genome exome resequencing of Zak and Zak*ERA8* were designed using PolyMarker with default settings (Table S4; Ramirez-Gonzalez et al. 2015b). The KASP assay for the naturally occuring preharvest sprouting susceptible (A660) versus resistant (C660) alleles of *TaMKK3-A* was performed as described in (Shorinola et al. 2017). Genomic DNA was extracted from BC_3_F_2_ leaf tissue on the Qiagen BioSprint-96, and DNA concentrations determined using the BioTek Gen5 plate reader. A standard 10 µL KASP reaction volume contained 15 ng/µL gDNA, 1x KASP Master Mix (LGC Genomics; V4.0 2x Mastermix 96/384; Low Rox), 0.42 µM reverse primer, and 0.168 µM each of allele 1 and 2 forward primers (Smith and Maughan 2015). KASP assays were performed on a thermocycler as follows: 1 cycle of 94 °C for 15 min, 10 cycles of 94 °C for 20 sec and 61-55 °C for 60 sec (starting at 61 °C, with 0.6 °C decrease in temperature per cycle), then 26 cycles of 94 °C for 20 sec and 57 °C for 60 sec (Biometra Tadvanced, Analytik Jena). FAM and HEX fluorescence was measured on the LightCycler 480 System (Roche).

In a preliminary experiment, 2,381 Zak/Zak*ERA8* BC_3_F_2_ lines were scored for WT versus ABA hypersensitive *ERA8* phenotype by plating on 2, 5, or 10 μM (+/-)-ABA. KASP assays were used to identify those lines containing recombination events within the *ERA8-*linked region. Of these, 424 were advanced to the BC_3_F_3_ and seeds were progeny tested for ABA sensitivity (Table S1). Based on Chi-squared analysis, BC_3_F_2:3_ from X5.1 (122 lines) and X5.2 (302 lines) showed Mendelian segregation for a single semi-dominant (or additive) gene (Fig. S1; Table S5). A linkage map of the X5.1 and X5.2 Zak/Zak*ERA8* BC_3_F_2:3_ populations was constructed using the regression algorithm and Kosambi mapping function in JoinMap v4.0 (Kosambi 1943; Van Ooijen 2006).

### Genotyping-by-sequencing

Genotyping-by-sequencing (GBS) of Louise/Zak*ERA8* RILs was performed on a Thermo-Fisher Scientific Ion-Proton system (Poland et al. 2012; Truong et al. 2012). Briefly, to construct GBS libraries, DNA was digested with *Pst*I and *Msp*I, then ligated to barcode adapters using T4 DNA ligase. Samples were purified, pooled, and sequenced. The USDA-ARS Western Regional Small Grains Genotyping Lab SNP marker pipeline was used to analyze the sequencing reads. This pipeline used built-in Linux commands and routines written in the Perl 5 programming language (http://www.perl.org) to remove extraneous bases 5’ to the *Pst*I site, trim the sequence reads to uniform length “tags” of 70 or 100 bases, then identify tags unique to each of the parent lines. These tags were aligned to the RefSeq v1.0 wheat genome (IWGSC et al. 2018) using SOAP2 allowing 0-2 mismatches (Li et al. 2009b). Tags that aligned to two or more chromosomes were discarded, leaving only tags that aligned to 0 (unknown) or 1 chromosome. A Perl 5 routine was then used to identify pairs of these filtered tags that differed by one or two SNPs between the parents, Louise and Zak*ERA8*, possibly representing alternate alleles. Linux built-in commands were then used to score the presence of those markers in the progeny population and a Perl 5 routine was used to convert the presence/absence data to genotype calls. Markers that were scored as missing (neither allele present) in more than 75 % of the progeny were removed from the data set. An iterative imputation of missing data was calculated using the ‘missforest’ package in R (Stekhoven and Bühlmann 2012). If a marker appeared to be heterozygous for 21 or more of the 207 RILs, then that marker was excluded from analysis to reduce false positive results.

### Genetic linkage map and QTL mapping

A linkage map of the Louise/Zak*ERA8* RIL population was constructed using SNP data from GBS using the maximum likelihood algorithm in JoinMap v4.0 (Van Ooijen 2006). Linkage groups of 5 or more markers were selected with a LOD above 7.0 (Table S6). The 92 of 2,234 markers showing segregation distortion, *p* < 2e-05, may have resulted from genotyping errors and so were omitted from the analysis. Composite interval mapping (CIM) was performed with the R/qtl package in R v3.2.5 using the Haley-Knott Regression algorithm, and the Kosambi mapping function with step = 0 (Haley and Knott 1992; Broman et al. 2003). Significant LOD scores were determined using the ‘scanone’ function in R/qtl with the Haley-Knotts method and 1,000 permutations for each trait. QTL graphs were constructed using the ‘ggplot2’ package v3.1.0 in R v3.2.5 (Wickham 2009).

Linkage analysis was performed in the Louise/Zak*ERA8* RIL, Zak/Zak*ERA8* BC_3_F_2_:F_3_, and Otis/Zak*ERA8* F_2_:F_3_ populations using genotype information from the *ERA8* EMS-induced KASP markers (Table S4; Supplementary Material 3). QTL analysis for both populations was conducted as described above for the Louise/Zak*ERA8* GBS data. The QTL analysis R script can be found in the public repository ‘ERA8-Mapping-2019’ on GitHub (https://github.com/shantel-martinez/ERA8-Mapping-2019).

### RNA sequencing

An RNA-seq experiment (2.5 million reads) was performed on the original BC_3_ parents Zak (Zak2.7.3.7.2.3.10.3) and Zak*ERA8* (BC_2_F_3_; 95.6.85.4). The RNA was purified from dark-imbibed seed tissue, and should not contain many transcripts involved with photosynthesis. Seeds were imbibed on 5 mM MES, pH 5.5 for 8 hours. Embryos were dissected and frozen in liquid nitrogen under a green safelight to avoid induction of photosynthesis genes. RNA was extracted from 20 dissected embryos per sample using a modified version of an RNA extraction technique which yielded an average of 2,300 ng/μL of RNA with a RIN > 8.0 (Onate-Sánchez and Vicente-Carbajosa 2008; Nelson and Steber 2017). This was replicated three times for each parent sample. Sequencing was performed using the Illumina HiSeq 2500 (WSU Genomics Core). Quality of the RNA reads were checked using FastQC and all samples had a quality score > 32 (Babraham Bioinformatics 2012). Pseudo-alignment and quantification of RNA-seq reads was conducted with kallisto-0.42.3 using the quant command with options -b 100 and -t 8 (Bray et al. 2016). The wheat RefSeq v1.0 transcriptome assembly (IWGSC et al. 2018) was used as the reference genome for this alignment. SAMtools was used to sort and index the BAM files v0.2.19 (Li et al. 2009a). Gene differential expression analysis was done using the ‘sleuth’ package in R (Pimentel et al. 2017).

## Results

### *ERA8* mapping by bulked segregant analysis of a backcross population

*ERA8* was mapped in the Zak/Zak*ERA8* backcross population relative to DNA polymorphisms induced by the EMS-mutagenesis of Zak (Fig. 1; Fig. S1). *ERA8* segregates as a single semi-dominant Mendelian trait in backcross populations when the phenotype is examined at optimized ABA concentrations and after-ripening time points (Schramm et al. 2013; Martinez et al. 2014). Consistent with this, the Zak/Zak*ERA8* BC_3_F_2_ population (X5.2) showed segregation as a single semi-dominant gene on 2 μM ABA at 5 weeks of after-ripening (χ^2^ = 1.62, p-value = 0.20; Table S2). The germination of the 300 BC_3_F_2:3_ lines exhibited a normal distribution in the presence of 2 μM ABA, and the corresponding, Zak WT and *ERA8* controls showed an average of 66.9 % (SD ± 16.3) and 16.8 % (SD ± 8.8) germination after 5 days of imbibition, respectively (Fig. 1b). Thus, an F_2:3_ line was considered to be homozygous WT (+/+) if the percent germination was above 73 % and to be homozygous *ERA8* (*ERA8*/*ERA8*) if below 24 %. These individuals were tested on ABA two more times to confirm the phenotype. The 15 BC_3_F_2:3_ lines that consistently showed high percent germination on ABA were defined as WT-like (WT_b:15) and the 17 individuals that consistently showed low percent germination on ABA were defined as *ERA8*-like (ERA8_b:17) (Fig. 1b).

**Fig. 1.**
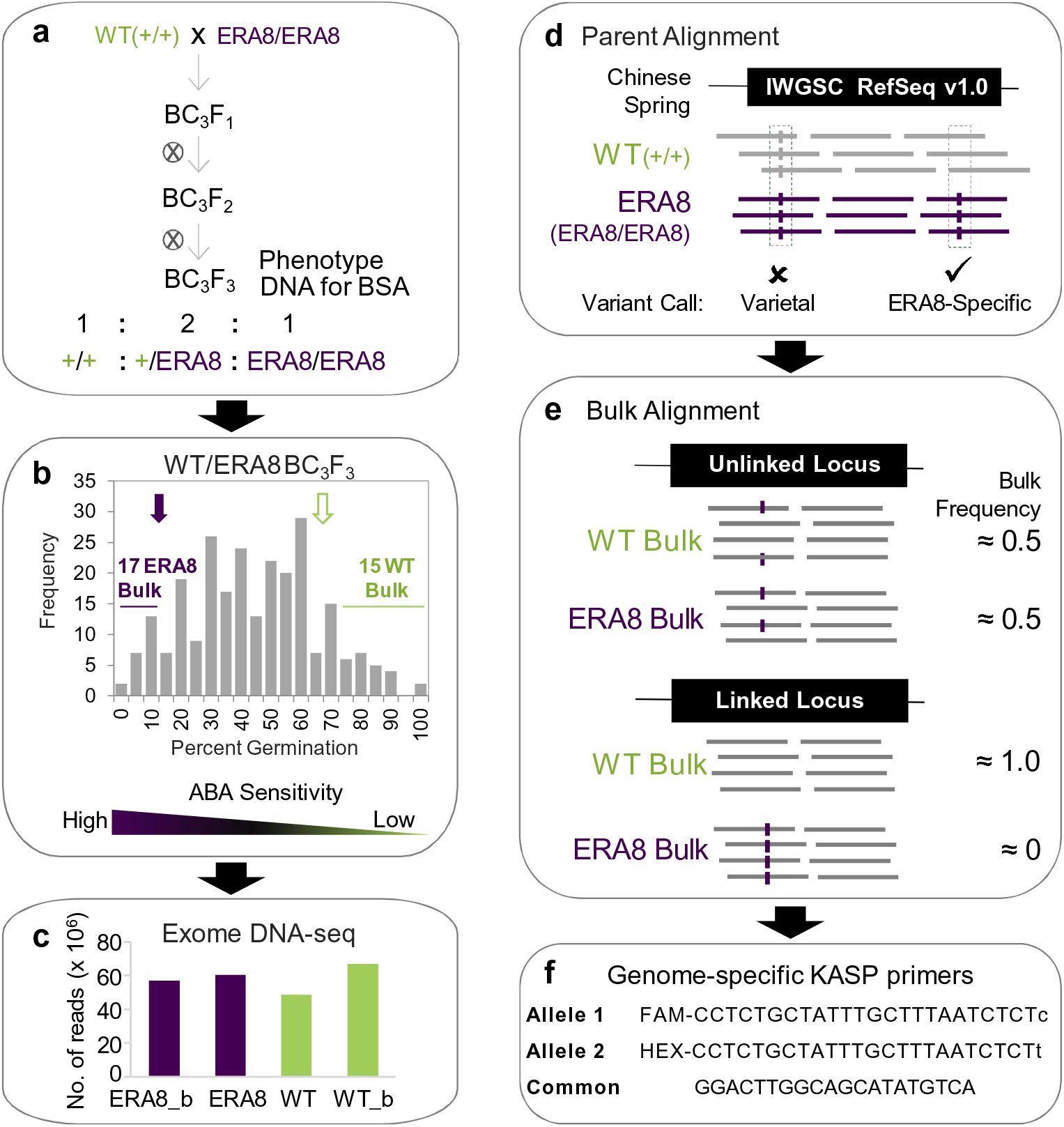
The BSA-exome-seq method. **a**) Genetics. The BC_3_F_2:3_ population was developed by crossing Zak WT (+/+) and BC_2_F_3_ Zak*ERA8* (*ERA8/ERA8*). BC_3_F_2:3_ genomic DNA was used for exome-seq. The phenotype was BC_3_F_2:3_ grain germination on ABA. **b**) Selection of bulked lines. Compared to the parental *ERA8* (purple arrow) and WT (green arrow) germination on ABA, 17 *ERA8*-like and 15 WT-like lines were chosen for bulked segregant analysis. **c**) The number of 125 bp paired-end Illumina HiSeq 2000 reads generated for each parent, WT bulk (WT_b), and *ERA8* bulk (*ERA8*_b) are shown. **d**) Alignment of parental reads to the Chinese Spring IWGSC reference sequence (v 1.0), where varietal differences from Chinese Spring were shared by the Zak WT and *ERA8* parents, and mutagen-induced (C/G to T/A) were *ERA8*-specific. **e)** Unlinked SNPs showed random variation in WT_b versus ERA8_b, whereas *ERA8*-linked SNPs did not. An example of an exon alignment and bulk frequency ratios are shown. **f)** KASP markers were designed to score SNPs. SNP_17 is shown.

Bulked segregant analysis was performed using bulked genomic DNA from the *ERA8*-like (*ERA8*_b) and WT-like (WT_b) BC_3_F_2:3_ lines. In order to maximize the number of informative sequences, genomic DNA was enriched for coding sequence using exome capture before performing high-throughput DNA sequencing on the Illumina HiSeq 2500 (48-67 million reads per sample; Fig. 1c; Table S3). Likely EMS-induced C to T and G to A polymorphisms between Zak WT and *ERA8* were identified by aligning to the RefSeq v1.0 wheat genome (Fig. 1d; IWGSC et al. 2018). SNPs identified between WT and *ERA8* were then screened against the *ERA8*-like and WT-like bulked exome sequences. If a region of the chromosome was not associated with the ABA-hypersensitive phenotype, then a polymorphic SNP should segregate randomly, appearing in about 50 % of the WT-bulk reads and 50 % of the *ERA8*-bulk reads (Fig. 1e). Conversely, if a SNP is associated with the ABA-hypersensitive phenotype, then the SNP allele from the *ERA8* parent should be found in nearly 100 % of the reads from the *ERA8* bulk and nearly 0 % of the WT-bulk reads. A large region of chromosome 4A showed strong enrichment for 80 EMS-induced mutations in the *ERA8*-like bulk sequences (Fig. 2). These 80 SNPs were in 46 genes spanning chromosome 4A (33 to 613 Mb). Of these, 42 SNPs in the coding region of 27 genes were located on the long arm of chromosome 4A. To our knowledge, this is the first example where an EMS-generated allele has been successfully mapped in allohexaploid wheat using bulked segregant analysis of high throughput DNA exome-seq data.

**Fig. 2.**
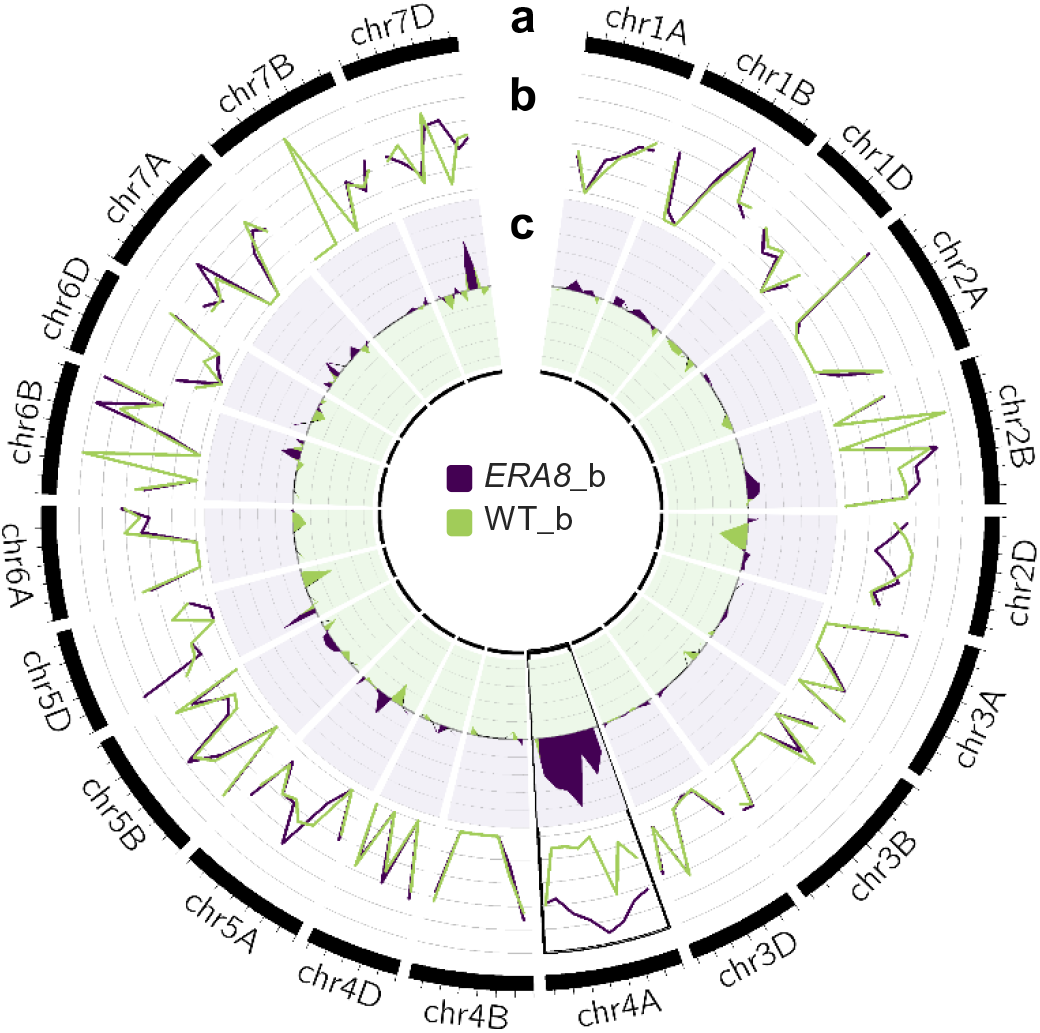
Bulk sequence frequencies aligned to the wheat genome. **a**) The outer track is the physical size (million bp per tick mark) of each chromosome based on the RefSeq v1.0 wheat reference genome sequence. **b**) Bulk frequencies (BF) of the *ERA8*_b (purple) and WT_b (green) were aligned to the reference genome and were calculated as a fraction of the alternate allele read depth over the total number of reads. **c**) The BF differences (BFD) were calculated from the differences between the BF of *ERA8*_b and the BF of WT_b.

### Fine mapping of *ERA8* in backcross and RIL populations

The *ERA8* map position was further refined using recombination events from a larger backcross population mapped relative to mutagen-induced SNPs. SNPs identified in the BSA-exome-seq analysis were scored by KASP assay (SNP_1 through SNP_30; Table S4). Based on an additional 424 random Zak/Zak*ERA8* BC_3_F_2:3_ lines, the *ERA8* phenotype was not associated with the centromere-proximal SNP_1 to SNP_6 region, but rather was strongly associated with a more distal region on the long arm of chromosome 4A spanning SNP_20, SNP_17, and SNP_29 (Fig. 3a). SNP_20 and SNP_29 flank a 4.6 Mb interval containing 70 high confidence predicted genes in the IWGSC RefSeq v1.0 annotation (Table S7). SNP_17, located within this interval, had the highest LOD score, ranging from 5.2. to 16.5 (Fig. 3a). The X5.2 population likely segregated for a weaker *ERA8* phenotype because it lost more dormancy due to longer storage in the freezer before plating (Table S1). Interestingly, SNP_17 is an *ERA8*-linked missense mutation in the cloned preharvest sprouting tolerance gene, *TaMKK3-A* (Nakamura et al. 2016; Torada et al. 2016; Shorinola et al. 2017). Because the *TaMKK3-A* gene was not present in the assigned chromosomes of the RefSeq v1.0 wheat genome reference assembly, its relative position was estimated based on neighboring genes in the draft WGA v0.4 sequence (IWGSC et al. 2014; Shorinola et al. 2017).

**Fig. 3.**
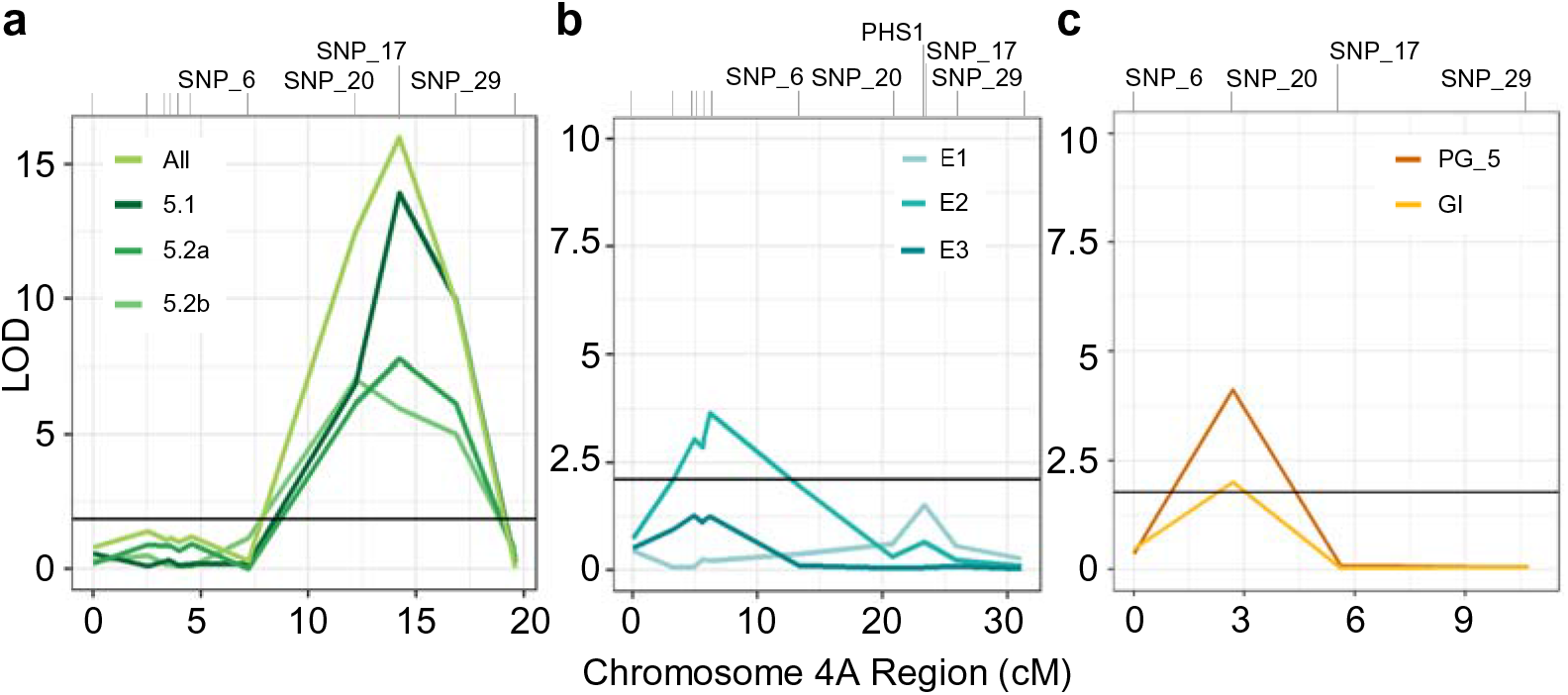
Fine mapping *ERA8* on chromosome 4A. QTL analysis was performed using mutagen-induced SNPs and germination index on ABA in the: **a**) Zak/Zak*ERA8* BC_3_F_3_ from crosses X5.1 and X5.2, **b**) Louise/Zak*ERA8* F_6:7_, and **c**) the Otis/Zak*ERA8* F_2:3_ populations. Population sizes were 424, 207, and 108 individuals, respectively. The Louise/Zak*ERA8* population was phenotyped over three environments (E1, E2, and E3). Marker distances (cM) of the 4A region of interest were calculated using JoinMap. SNP positions are marked above the graph.

Based on the fine mapping, *ERA8* appears to be within the 4,597,054 bp region between SNP_20 and SNP_29 (Fig. 3a; Fig. 4a-c). This region contains 70 genes of which 3 carry EMS-induced mutations including SNP_20 in a predicted GSK1 transcription factor (*TraesCS4A01G312200*), SNP_17 in the *TaMKK3-A* gene (*TraesCSU01G167000*) and SNP_29 in a gibberellin 20-oxidase gene (*TraesCS4A01G319100*). Two additional SNPs were found in low-confidence genes in the interval: SNP_33 in a suspected pseudogene, and SNP_34 in a low confidence predicted Ser/Thr phosphatase (Table S7). Unfortunately, we were unable to develop KASP markers to clearly distinguish between alleles of SNP_33 and SNP_34. The exome-seq analysis selected only C to T and G to A transitions for mapping in order to filter out sequence differences that were not caused by EMS mutagenesis. Additional analysis detected no insertion or deletion (in/del) polymorphisms within the SNP_20 to SNP_29 interval (Fig. S3). This in/del analysis detected a large deletion on the distal arm of chromosome 2DL and a rearrangement between chromosome arms 6AL and 6BS in *ERA8*. However, neither of these polymorphisms were linked with increased ABA sensitivity (Fig. 2).

A comparison of phenotype to the location of recombination events was used to further define the *ERA8* position relative to the four mutagen-induced SNPs in the *ERA8*-linked region (SNPs 6, 20, 17 and 29; Fig. 4; Table S4). Homozygous lines were placed into haplotype groups (I to VII) based on the location of recombination events (Fig. 4d-e). Then the germination phenotype of each group was placed into the *ERA8*-like (purple) or WT-like (green) class based on whether or not they showed a statistically significant difference from the *ERA8* or Zak WT parent based on Student’s t-test. The location of recombination events (Fig. 4d) was compared to the germination phenotype on ABA (Fig. 4e). Based on the germination phenotypes of Group I and Group IV, SNP_29 does not appear to determine the *ERA8* phenotype. Based on Groups III, IV, and VI, the *ERA8*-determining mutation is most tightly linked to the SNP_20 to SNP_17 region containing 25 predicted genes. Based on Groups II, V, and VI, the region from SNP_6 to SNP_20 is not sufficient to determine the germination phenotype, although this conclusion is supported by fewer lines. All haplotypes carrying SNP_17 have an *ERA8* phenotype, suggesting that either SNP_17 or a tightly linked non-coding region mutation between SNP_20 and SNP_29 cause the *ERA8* phenotype. Unfortunately, our ability to fine-map *ERA8* is severely limited by having only the four mutagen-induced SNPs in the *ERA8* region.

**Fig. 4.**
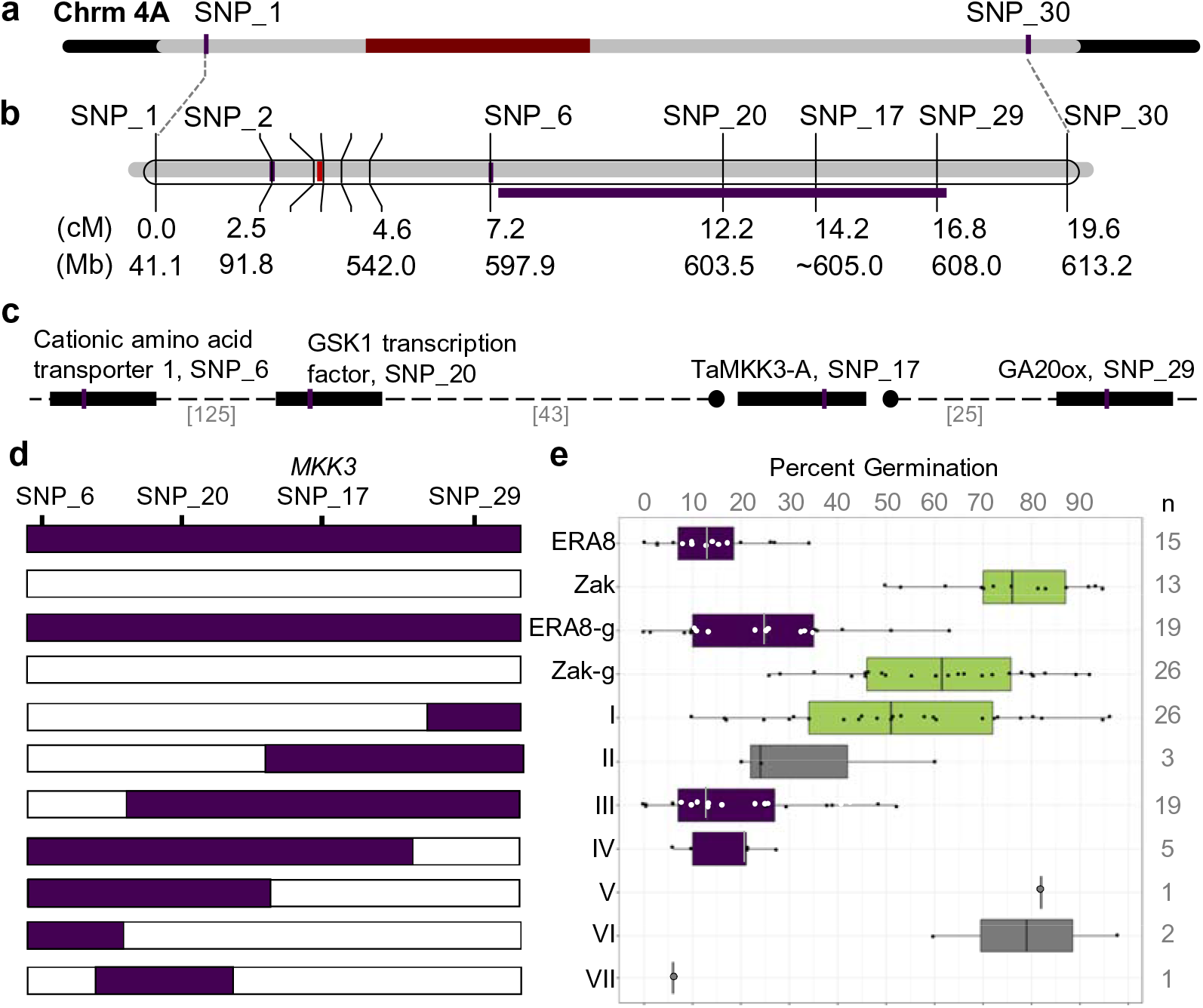
The *ERA8* interval. **a**) Initial BSA based on exome-seq of 32 BC_3_F_2:3_ identified the *ERA8* region (grey) on chromosome 4A. The centromeric region is in red. **b**) QTL analysis relative to EMS-induced SNPs in 424 BC_3_F_2:3_ mapped a 4.6 cM region containing the *ERA8*-linked interval (purple bar) spanning **c**) the 4 unique genes containing SNP_6 through SNP_29. The number of predicted genes between SNPs is in brackets. The *TaMKK3-A* position is estimated based on previous mapping studies. **d)** Lines containing recombination events in the SNP_6 to SNP_29 region were placed into groups I to VII based on whether they carried the *ERA8* (purple) or Zak WT (white) SNP allele. **e**) The ABA germination phenotype of groups I-VII is shown as a box and whisker plot where dots show the percent germination of individual BC_3_F_2:3-4_ lines. The population parents *ERA8* and Zak, and BC_3_F_2:3_ line where all four SNPs had the *ERA8* genotype (*ERA8*-g) or Zak genotype (Zak-g) are shown as controls. *ERA8*-like (purple) and Zak-like 21 (green) groups were determined based on Student’s t-test (p<u>≤</u>0.05) compared to the parents. n = the number of lines per group.

### *ERA8* mapping by QTL analysis of a recombinant inbred line population

A Louise/Zak*ERA8* RIL population of 225 lines was created to confirm the *ERA8* map position by QTL analysis of the ABA hypersensitive germination phenotype. QTL analysis was performed using germination data from three environments referred to as E1, E2, and E3 (Table S8). Because dormancy and wheat ABA sensitivity vary with environment, the RIL plating assays were performed using the after-ripening time point and ABA concentration resulting in the strongest germination difference between the Louise and Zak*ERA8* parents (Fig. 5a). It proved difficult to find conditions that differentiated between Louise and Zak*ERA8* because Louise had more ABA sensitivity than the original wild-type Zak parent. When whole-seed germination was examined, *ERA8* was only slightly more ABA sensitive than Louise (5.6 % versus 25.6 % germination at 6 weeks of after-ripening). A stronger difference between Louise and Zak*ERA8* was observed when grains were cut to break coat-imposed dormancy, and ABA sensitivity examined during embryo germination (Fig. 5b). For example, in E1 with 7 weeks of after-ripening, Louise and *ERA8* showed 92 % and 4 % germination on 5 µM ABA, respectively. In E1, the strongest difference in Louise and *ERA8* germination was observed at day one of imbibition (Table S8). Both percent germination and GI were used in the QTL analysis because the percent germination of the RILs had a normal distribution, whereas the GI distribution was skewed towards the non-dormant parent Louise (Table S8; Fig. 5c). While there was a stronger difference between parents in E2 and E3 using germination index (GI) than using percent germination, all of the significant QTL were identified using percent germination data.

**Fig. 5.**
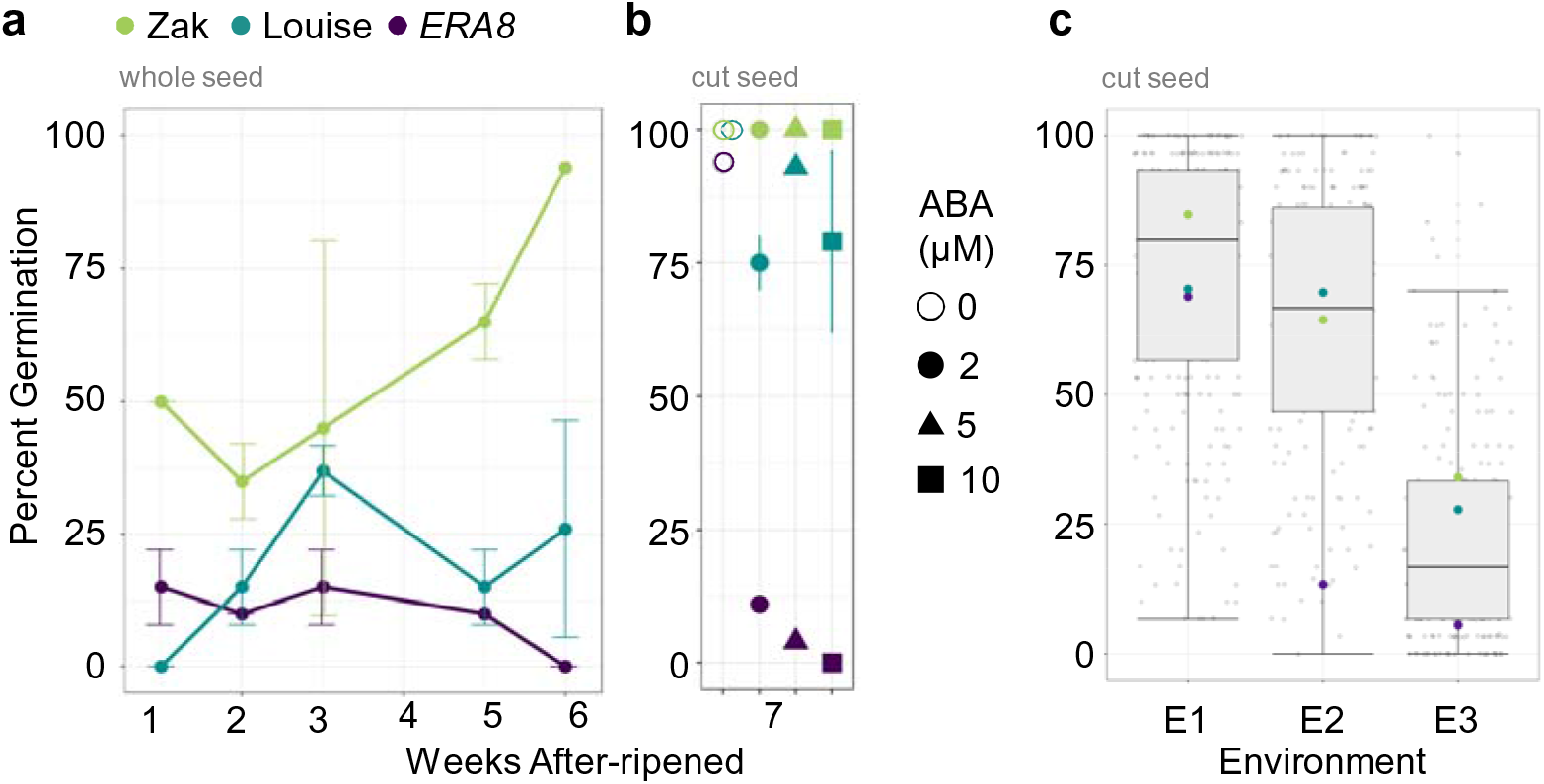
ABA sensitivity of Louise (blue), *ERA8* (purple), Zak WT (green), and the Louise/Zak*ERA8* RIL(grey) population. The parents (Louise, *ERA8*, and Zak WT) percent germination after 5 days of imbibition over an after-ripening time course in environment 1 were assayed in: **a**) whole seeds on 2 µM ABA and **b**) cut seeds on 0, 2, 5, and 10 µM ABA. **c**) The RIL population was assayed across three environments with the percent germination after 2, 3, and 1 day of imbibition on 5µM, 2µM, and 2µM of ABA shown, respectively. Environments: E1) F_6_ seed from the greenhouse 2013; E2) F_7_ seed from the field 2014; and E3) F_7_ seed from the field 2015

Both Louise and *ERA8* contributed multiple QTL impacting ABA sensitivity during germination. Linkage analysis of GBS data identified 2,234 single nucleotide polymorphisms (SNPs) between Louise and *ERA8* that fell into 45 linkage groups spanning all 21 chromosomes (Table S6). Seven significant QTL mapped to chromosomes 2D, 4A, 4B, and 7B (Table 1). None of the QTL were significant in multiple environments. Louise contributed higher ABA sensitivity due to the *QABA.wsu-2D* and *QABA.wsu-4A.3* loci. In fact, *QABA.wsu-2D* had the highest LOD score of 5.69. *ERA8* contributed higher ABA sensitivity due to *QABA.wsu-4A.1*, *QABA.wsu-4A.2*, *QABA.wsu-4B*, *QABA.wsu-7B.1,* and *QABA.wsu-7B.2* loci. The two major *ERA8*-associated loci were on chromosome 4A, with LOD scores of 5.32 and 5.05, respectively. None of these QTL was associated with a strong ABA hypersensitive phenotype because they were significant with only one to three days of imbibition. This suggests that Louise and Zak*ERA8* did not differ greatly for ABA sensitivity. The QTL identified, however, had additive effects since RILs carrying an increasing number of ABA sensitivity loci had increasing ABA sensitivity (Fig. S4). None of the QTL identified co-localized with QTL for plant height mapped in the same RIL population (Fig. S5; Table S9). However, a Louise/Zak*ERA8* heading date QTL co-localized with *QABA.wsu-4A.9* in the combined analysis using GBS and EMS-generated SNPs.

To determine whether either of the *ERA8* QTLs on chromosome 4A co-localized with the *ERA8* region mapped in the backcross population, QTL analysis of the Louise/Zak*ERA8* RILs was performed using only KASP markers for 12 mutagen-induced SNPs. QTL mapping identified one significant QTL on chromosome 4A, located at the 4A centromere (*QABA.wsu-4A.2,* LOD 3.5) (Fig. 3b). Thus, the *ERA8* phenotype was not significantly associated with SNP_17 in the Louise/Zak*ERA8* RIL population. There are two possible explanations for this result, either there are other loci in the Louise background that suppress the *ERA8* phenotype, or both Louise and Zak*ERA8* contributed independent ABA-sensitive alleles of *TaMKK3-A* to the RIL population.

### Alleles of *TaMKK3-A*

Based on the hypothesis that the *ERA8* phenotype may have resulted from a mutation in the *TaMKK3-A* coding region, we characterized the *TaMKK3-A* alleles of Zak, Zak*ERA8*, and Louise using KASP assays (Shorinola et al. 2017). The naturally-occuring PHS tolerant allele of *TaMKK3-A/Phs1* in wheat results from a C nucleotide at position +660 associated with a recessive dormant phenotype, whereas the PHS susceptible allele is an A nucleotide at position +660 (Torada et al. 2016). Interestingly, KASP assays showed that Louise has the dormant *TaMKK3-A*-C660^R^ allele, whereas wild-type Zak and Zak*ERA8* carry the susceptible *TaMKK3-A*-A660^S^ allele (Fig. 6; Fig. S6; Shorinola et al., 2017). Only the Zak*ERA8* line carries the EMS-induced G to A transition at position +1093 of *TaMKK3-A*. This missense mutation results in a change from glutamic acid (E) to the basic amino acid lysine (K) at position 365 within the conserved Nuclear Transport Factor 2 (NTF2) domain. Thus, Zak*ERA8* and Louise carry two different *MKK3* alleles associated with higher dormancy and ABA sensitivity. This suggests that the dormant *ERA8* phenotype failed to segregate in the Louise/Zak*ERA8* RIL population because both parents contributed dormant alleles of *MKK3*. Thus, the QTL mapped were due to contributions from other loci affecting ABA sensitivity in the population. If this hypothesis is correct, then the ABA hypersensitive phenotype should map to the *ERA8* region identified in the backcross population if Zak*ERA8* is crossed to a line carrying the nondormant *TaMKK3-A*-A660^S^ allele, such as hard white Otis. QTL analysis of 108 Otis/Zak*ERA8* F_2_:F_3_ lines detected a significant QTL associated with ABA-hypersensitive germination at SNP_20, suggesting that *ERA8* is in the SNP_20 to SNP_29 region (Fig. 3c). Taken together, these results support the hypothesis that the *ERA8* phenotype is due to the EMS-induced *TaMKK3-A-*A1093^R^ allele.

**Fig. 6.**
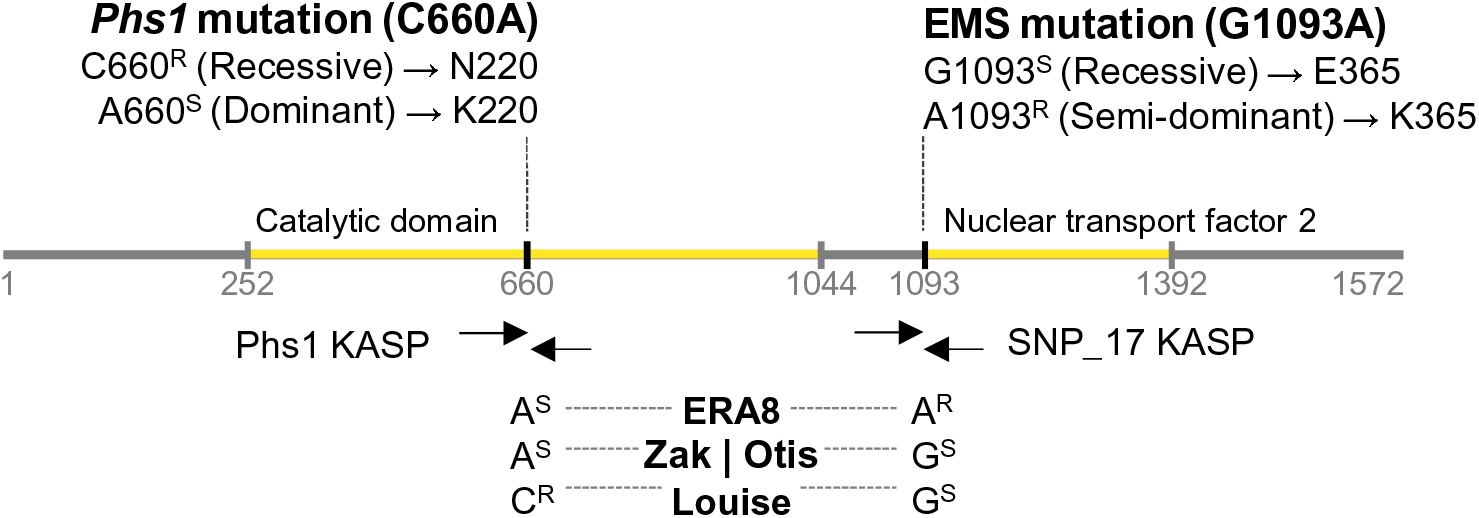
Alleles of the *TaMKK3-A* gene. Increased dormancy and PHS tolerance may result either from the recessive *Phs1* allele at +660 bp or from the semi-dominant EMS-induced *ERA8* allele at +1093 bp of *MKK3-A*. Nucleotide and amino acid differences are shown for the S (susceptible) and R (resistant) alleles. The genotypes of *ERA8*, Zak, Otis, and Louise are shown at both positions.

### Differentially expressed genes in *ERA8*

There are 70 genes in the SNP_20 to SNP_29 region. If any gene in this region resulted in an ABA-hypersensitive phenotype due to non-coding region mutations such as a promoter or intron, then we would expect to see a difference in expression levels in *ERA8* versus wild-type Zak. The semi-dominant germination phenotype would then result from a change in the dosage of a regulatory gene. RNA-seq was used to identify additional *ERA8* candidate genes in the *ERA8*-linked region based on differential expression in imbibing Zak*ERA8* versus wild-type seeds. Differences in seed dormancy should result in many transcriptome differences downstream of the *ERA8* mutation (Barrero et al. 2009; Liu et al. 2013a; Nelson and Steber 2017). To focus on transcriptome differences possibly resulting from an *ERA8* promoter or intron mutation, transcriptome analysis was performed comparing *ERA8* and Zak WT seed after-ripened long enough to reduce differences in seed dormancy. Zak*ERA8* BC_2_F_3_ and Zak WT seeds after-ripened for 1.5 years both showed 100% germination within two days of imbibition. The mRNA was prepared from seeds imbibed for 8 h without ABA in the dark. Embryos were harvested into liquid nitrogen under a green safe light to enrich for transcripts expressed during seed germination while avoiding the induction of highly expressed photosynthesis-related transcripts that can represent as much as 50% of RNA-seq reads from photosynthetic tissue (Trick et al. 2012). The most significantly differentially-regulated genes were located on chromosomes 2B (*ERA8*-up/WT-down) and 2D (WT-up/*ERA8*-down) (Table S10). On chromosome 4A, there were four WT-up/*ERA8*-down genes, and eight *ERA8*-up/WT-down genes (Table S10). Only two differentially-expressed genes were within the SNP_6 to SNP_30 interval, both homologues of BED Zinc finger family proteins believed to be involved in stress response (Joly-Lopez et al. 2017). However, both BED Zinc finger homologues were located between SNP_29 and 30, indicating that they are not in the strongest *ERA8*-linked region between SNP_ 20 and SNP_29. *TaMKK3-A* did not show any significant change in expression in WT versus *ERA8* under the conditions examined. In conclusion, the RNA-seq data did not reveal additional *ERA8* candidate genes based on differential expression analysis from the time point examined.

## Discussion

### BSA-exome-seq as a method to map EMS-induced mutations

Here we combine two previously published methods, exome capture and bulked segregant analysis of a backcross population, to map a mutagen-induced trait with variable expressivity in the large allohexaploid wheat genome. Exome capture was previously implemented to characterize all of the coding-region EMS-induced mutations in a wheat TILLING population (Krasileva et al. 2017). BSA was previously used to map genes relative to varietal DNA polymorphisms identified by RNA-seq (Trick et al. 2012). We combined these two approaches to map a mutation using bulked segregant analysis of exome sequence from pools of genomic DNA from WT-like and *ERA8*-like individuals from a BC_3_F_2:3_ population. A similar approach was used in the tetraploid Kronos TILLING population to map tall versus short EMS-induced alleles from a segregating M_4_ population and a leaf chlorosis phenotype in an F_2_ population (Mo et al. 2017; Harrington et al. 2019). In both of these studies, the phenotypic differences between wild-type and mutant individuals were relatively large compared to the more difficult-to-phenotype *ERA8* mutation. BSA of a backcross population was particularly useful for mapping *ERA8* since variation in dormancy is a quantitative trait in RIL populations. BSA allowed the use of a relatively small number of confidently phenotyped BC_3_F_2_:F_3_ individuals. It was beneficial to map in an *ERA8* BC_3_ population because the mutant plants became healthier and the ABA-hypersensitive phenotype more consistent with additional backcrosses. This may have occurred because background EMS mutations were removed by backcrossing (Uauy et al. 2017). Researchers interested in mapping mutant phenotypes with incomplete penetrance, variable expressivity, or requiring careful developmental staging may find it helpful to consider additional backcrosses before investing in BSA-exome-seq.

### Comparison of *ERA8* mapping by QTL analysis and BSA-exome-seq

There are both advantages and disadvantages to mapping in an RIL population as opposed to a backcross population. Creation of RIL populations may require more time than creating a population from a single backcross event since plants must be advanced by single seed descent to at least the F_5_ generation. Moreover, mapping in a backcross population simplifies the genetics, decreasing the impact of other loci. BSA of the backcross population had the advantage that the ABA-hypersensitive phenotype showed Mendelian segregation because *ERA8* was the major locus contributing the ABA-hypersensitive phenotype (Table S2; Table S5; Martinez et al. 2014). While Louise was selected as an RIL population parent based on having dormancy characteristics similar to Zak, we were blind to the precise genetic alleles contributing to the dormancy or lack of dormancy in Louise (Fig. 5). Thus, the fact that Louise carried a dormant allele of the *TaMKK3-A* gene was a surprise, making Louise less than an ideal choice given the hypothesis that *ERA8* is a new dormant allele of *TaMKK3-A*. Both the Louise and Zak*ERA8* parents contributed other QTL associated with higher ABA sensitivity during germination to the RIL population (Table 1; Table S9). These QTL are of potential use for PHS tolerance breeding in northwestern U.S. wheat since pyramiding of multiple ABA-sensitive QTL provided increasing seed dormancy (Fig. S4). One drawback, however, to the mapping in the backcross instead of RIL population was the paucity of polymorphisms in the *ERA8* interval. We were unable to confidently localize *ERA8* to a region smaller than 4,597,054 bp due to a lack of additional polymorphisms between SNP_20 and SNP_29 (Table S7). A mapping population derived from a cross between near-isogenic lines containing varietal differences within one chromosome segment may have the advantage of having a higher density of polymorphisms (Yan et al. 2003; Trick et al. 2012). But such a population has the drawback that there may be multiple sequence polymorphisms within a candidate gene(s), making it more challenging to identify the mutation causing the phenotype of interest (Shorinola et al. 2017). Future efforts to determine if the *ERA8* phenotype is caused by the *TaMKK3-A-G1093A* allele could exploit additional polymorphisms identified by complete chromosome sequencing of the Zak*ERA8* mutant chromosome (as in Harrington et al. 2019), or make use of a population where both parents carry the PHS susceptible *TaMKK3-A-A660^S^* allele, such as a population derived from Otis/Zak*ERA8*.

### *TaMKK3-A* is the strongest *ERA8* candidate gene

Several lines of evidence indicate that *TaMKK3-A* is the strongest candidate for the *ERA8* gene. First, SNP_17/*TaMKK3-A*-G1093A is the strongest QTL mapped in the backcross population, with a LOD score as high as 16.5 (Fig. 3a). Based on examination of recombination events in the region, it does not appear that either SNP_20 or SNP_29 are required for the *ERA8* phenotype (Fig. 4d,e). Second, G1093A was the only EMS-induced mutation identified in a high confidence gene within the *ERA8*-linked interval between SNP_20 and SNP_29 (Table S7). The only other mutations in this interval were SNP_33 in a pseudo-gene and SNP_34 in a low confidence gene (*TraesCS4A01G473200LC*). Third, there were no genes in the SNP_20 to SNP_29 interval showing differential expression between Zak and Zak*ERA8* during seed imbibition, when *ERA8* is expected to be expressed (Table S10). Thus, it does not appear likely that the *ERA8* phenotype results from a promoter mutation in this interval leading to altered expression of a regulatory gene. Fourth, *ERA8* could be mapped to the region flanked by SNP_20 and SNP_29 in the Otis/Zak*ERA8* population, but not in Louise/Zak*ERA8* RIL population (Fig. 3b,c). This appears to be an inadvertent complementation test where both Louise and Zak*ERA8* carry dormant alleles and Otis a non-dormant allele of *TaMKK3-A*. Finally, *MKK3* is a strong candidate for *ERA8* because it is known to function in wheat and barley grain dormancy as well as in Arabidopsis ABA signaling (Danquah et al. 2015; Nakamura et al. 2016; Torada et al. 2016; Shorinola et al. 2017). Taken together, these data suggest that the G1093A mutation in *TaMKK3-A* is the most likely cause of the *ERA8* phenotype, although we cannot discard the presence of other mutations in the 70-gene SNP_20 to SNP_29 interval. Future work will need to determine if the *TaMKK3-A*-G1093A mutation results in an ABA hypersensitive germination phenotype when transformed into Zak or another variety carrying the PHS susceptible *TaMKK3-A-*A660^S^ allele.

### MKK3 protein family structure and function

MKK3 family predicted proteins contain a conserved ATP-binding domain, a kinase domain, and an NTF2 (Nuclear Transport Factor2)-like domain. Both of the natural dormant *MKK3* variants of wheat and barley are amino acid polymorphisms in the kinase domain, lysine (K) 220 to asparagine (N) in wheat and asparagine (N) 260 instead of threonine (T) in barley (Nakamura et al. 2016; Torada et al. 2016). Interestingly, the dormant wheat *MKK3-A-*C660^R^ allele is recessive whereas the dormant barley allele is dominant. Thus, it appears that in the *Triticeae* a dormant phenotype can result from either gain and loss-of-function alleles of *MKK3*. If *ERA8* is an allele of *TaMKK3-A*, then the fact that *ERA8* is semi-dominant suggests that MKK3 is a positive regulator of ABA signaling in wheat. Previous work, however, suggests that the recessive dormant *MKK3-A-*C660^R^ is the ancestral allele. In order to define whether *TaMKK3-A* is a positive or negative regulator, future work will need to determine whether a null allele results in increased or decreased seed dormancy and ABA sensitivity.

The semi-dominant *ERA8* phenotype is linked to the G1093A mutation resulting in a change from glutamate (E) 365 to lysine (K) within the conserved NTF2-like domain (Danquah et al. 2015). Since E365K is a change from a negatively to positively charged amino acid, this change may alter protein function. The NTF2 domain is unique to the *MKK3* family of map kinases (Jiang and Chu 2018) that appears to be involved in protein-protein interaction with downstream MKK3-activated group C MAP kinases, MPK7, 1, and 3 (Dóczi et al. 2007). A semi-dominant phenotype might result from increased affinity for one of the wheat MKK3-target proteins.

*TaMKK3-A* is expressed in many tissues where ABA signaling is expected to occur such as roots, leaves, seedlings, and grain (Torada et al. 2016; Ramírez-González et al. 2018). But the transcript levels are highest in maturing embryos, when dormancy and desiccation tolerance are established. The *ERA8* mutation alters ABA sensitivity in seeds but not during leaf stomatal closure (Schramm 2010). In contrast, the ABA hypersensitive *Warm4* (*Wheat ABA-responsive mutant 4*) mutant of wheat exhibited altered ABA sensitivity both during seed germination and in vegetative tissues (Schramm et al. 2010). This observation suggests that an *ERA8* mutation in the *MKK3* NTF2 domain may increase binding to a target protein involved in seed but not leaf ABA signaling.

### The potential role of *MKK3* signaling in controlling wheat grain dormancy and germination

If the *ERA8* phenotype truly results from the *TaMKK3-A*-G1093A mutation, then future work will need to further investigate its role in wheat ABA signaling and seed dormancy. The recovery of *MKK3* alleles providing increased dormancy and PHS tolerance in wheat and barley based both on natural variation and chemical mutagenesis would suggest that *MKK3* is a key regulator of these responses in cereals. This likely has broad relevance in plants since the *MKK3* gene family is highly conserved in eukaryotes, including Arabidopsis (*AtMKK3*), barley (*HvMKK3*), rice (*OsMKK3*), and tobacco (*NtNPK2*) (Shibata et al. 1995; Xiong and Yang 2003; Hamel et al. 2006; Nakamura et al. 2016; Torada et al. 2016). The mitogen-activated kinase signaling cascade was initially characterized based on its role in controlling mitosis and cell growth in yeast and humans (reviewed by Rodriguez et al. 2010). Identification of ABA-activated mitogen-activated protein kinases (MAPKs) in the barley aleurone system first suggested a role for MAPKs in ABA regulation of cereal grain germination (Knetsch et al. 1996). MAP kinases phosphorylate proteins in a wide range of signaling pathways. The Arabidopsis MAP kinase *MKK3* performs jasmonic acid (JA)-stimulated phosphorylation of the MAP kinase *MPK6*, which in turn negatively regulates the MYC2/JIN1 transcription factor, a positive regulator of root growth (Takahashi et al. 2007). This is interesting given that JA hormone appears to stimulate germination in barley and in wheat (Barrero et al. 2009; Tuttle et al. 2015; Xu et al. 2016; Martinez et al. 2016). *MKK3* and *MPK6* also have roles in ABA signaling and seed dormancy in Arabidopsis, together with other map kinases *MPK3, MPK17, MPK18*, *MKK1*, *MKK9, MPK1*, *MPK*2*, MPK7*, and *MPK14* (Liu 2012; Danquah et al. 2015). Loss of *AtMKK3* function in Arabidopsis led to ABA-hypersensitive seed germination (Danquah et al. 2015). Future work will need to identify TaMKK3-A target proteins, and to define domain functions through structure-function analyses. The *ERA8* allele could be a useful tool for analysis of protein-protein interactions in wheat. Such analyses would need to examine MKK3 domain functions in the context of ABA responses including seed germination, root elongation, and stomatal closure.

### Breeding for PHS tolerance with *ERA8*

Even if *ERA8* is not an allele of *TaMKK3*, the identification of SNP_17 as a strong *ERA8*-linked marker will have utility for deploying *ERA8* in breeding programs. Before crossing to *ERA8*, it will be important to determine if the starting *MKK3* allele is the susceptible A660 or the tolerant C660, in order to know which dormancy alleles will be segregating in the cross. Deploying an EMS-induced mutation is advantageous since the marker associated with the *ERA8* allele will be polymorphic in most outcrosses. The exome-seq data for *ERA8* revealed a large deletion on chromosome arm 2DL and an insertion on 6BS, likely induced by mutagenesis. To avoid possible negative effects of these in-dels, it would be wise to select against them during the breeding process. Since *ERA8* was selected in a soft white wheat background, this gene can be used to breed for PHS resistance regardless of seed coat color. Future breeding efforts will need to move *ERA8* into a hard wheat background to produce bread products versus soft wheat products. Previous work established that *ERA8* has sufficient dormancy to prevent preharvest sprouting at maturity, but loses dormancy rapidly enough through 8-10 weeks of after-ripening to allow fall planting of winter wheat without a strong negative impact on emergence or yield (Martinez et al. 2014, 2016). Farmers would need to be advised to wait 4-9 weeks before replanting *ERA8*-carrying winter wheat. *ERA8* has also been shown to maintain higher seed dormancy than Zak at temperatures as low as 20 °C (Martinez et al. 2016). Maintaining dormancy at lower temperatures is beneficial since cooler conditions exacerbate PHS.

## Acknowledgments

The authors thank Tracy Harris and Adrienne Burke for expert technical assistance, Nuan Wen for expert advice on KASP assays, and Yiyong (Ben) Liu for sharing his expertise and advice on Illumina RNA sequencing. Thanks are due to M. Pumphrey, K. Garland Campbell, and members of the Steber lab for assistance with planting, sampling, and harvesting. The authors also wish to thank the International Wheat Genome Sequencing Consortium (IWGSC) for pre-publication access to the wheat genome RefSeq v1.0. Special thanks go to K. Garland Campbell and to members of the Garland Campbell and Steber labs for helpful feedback on the research and manuscript. This work was funded by NIFA 2015-05798 (to CMS), the Washington Grain Alliance (to CMS), the USDA-ARS (to CMS), the Royal Society FLAIR Fellowship FLR_R1_191850 (to OS), and the Biotechnology and Biotechnology and Biological Sciences Research Council Designing Future Wheat Institute Strategic Programme BB/P016855/1 (to CU).

## Supplemental Material 1 (PDF 555 kb)

Fig. S1 - Crossing strategy to generate the *ERA8* backcross mapping population

Fig. S2 - WT and *ERA8* parental seed germination assay

Fig. S3 - Insertions or deletions between WT and *ERA8*

Fig. S4 - ABA response to number of QTL in the Louise/Zak*ERA8* RILs

Fig. S5 - QTL analysis of Louise/Zak*ERA8* heading date and height

Fig. S6 - WT and *ERA8* coding sequence of *TaMKK3-A*

## Supplemental Material 2 (PDF 424 kb)

Table S1 - Summary of Zak/Zak*ERA8* ABA phenotyping conditions

Table S2 - Chi-squared analysis of Zak/Zak*ERA8* BC_3_F_2:3_ X5.2

Table S3 - Exome capture sequencing quality and statistics

Table S4 - Primers created from the GBS and exome capture identified SNPs

Table S5 - Chi-squared analysis of Zak/Zak*ERA8* BC_3_F_2:3_ X5.1 and X5.2

Table S6 - Louise/Zak*ERA8* linkage group summary

Table S7 - Genes in the *ERA8* 4A region

Table S8 - Summary of Louise/Zak*ERA8* ABA phenotyping conditions

Table S9 - Significant QTL of ABA sensitivity, height, and heading date for Louise/Zak*ERA8*

Table S10 - Genes differentially expressed between Zak and *ERA8*

## Supplemental Material 3 (XLSX 1,671 kb)

Rqtl input files that include the genotypic, phenotypic, and map information for all mapping populations.

### Compliance with ethical standards

The authors declare that they have no conflict of interest.

## Data availability

The datasets generated during and/or analyzed during the current study are available in the GitHub repository, <u>ERA8-Mapping</u>.

## References

Allen AM, Barker GLA, Wilkinson P, et al (2013) Discovery and development of exome-based, co-dominant single nucleotide polymorphism markers in hexaploid wheat (*Triticum aestivum* L.). Plant Biotechnol J 11:279–295. doi: 10.1111/pbi.12009

Babraham Bioinformatics (2012) FastQC A Quality Control tool for High Throughput Sequence Data. http://www.bioinformatics.babraham.ac.uk/projects/fastqc/

Barrero JM, Talbot MJ, White RG, et al (2009) Anatomical and transcriptomic studies of the coleorhiza reveal the importance of this tissue in regulating dormancy in barley. Plant Physiol 150:1006–1021

Bewley JD, Bradford KJ, Hilhorst HWM, Nonogaki H (2013) Seeds. Springer New York, New York, NY

Bolger AM, Lohse M, Usadel B (2014) Trimmomatic: a flexible trimmer for Illumina sequence data. Bioinforma Oxf Engl 30:2114–2120. doi: 10.1093/bioinformatics/btu170

Bray NL, Pimentel H, Melsted P, Pachter L (2016) Near-optimal probabilistic RNA-seq quantification. Nat Biotechnol 34:525–527. doi: 10.1038/nbt.3519

Broman KW, Wu H, Sen Ś, Churchill GA (2003) R/qtl: QTL mapping in experimental crosses. Bioinformatics 19: 889–890. doi: 10.1093/bioinformatics/btg112

Chapman JA, Mascher M, Buluç A, et al (2015) A whole-genome shotgun approach for assembling and anchoring the hexaploid bread wheat genome. Genome Biol 16:26. doi: 10.1186/s13059-015-0582-8

Cingolani P, Platts A, Wang LL, et al (2012) A program for annotating and predicting the effects of single nucleotide polymorphisms, SnpEff. Fly (Austin) 6:80–92. doi: 10.4161/fly.19695

Danquah A, de Zélicourt A, Boudsocq M, et al (2015) Identification and characterization of an ABA-activated MAP kinase cascade in *Arabidopsis thaliana*. Plant J 82:232–244. doi: 10.1111/tpj.12808

DePauw RM, McCaig TN (1991) Components of variation, heritabilities and correlations for indices of sprouting tolerance and seed dormancy in *Triticum* spp. Euphytica 52:221–229. doi: 10.1007/BF00029399

Dóczi R, Brader G, Pettkó-Szandtner A, et al (2007) The Arabidopsis Mitogen-Activated Protein Kinase Kinase MKK3 Is Upstream of Group C Mitogen-Activated Protein Kinases and Participates in Pathogen Signaling. Plant Cell 19:3266–3279. doi: 10.1105/tpc.106.050039

Finkelstein RR, Reeves W, Ariizumi T, Steber CM (2008) Molecular aspects of seed dormancy. Annu Rev Plant Biol 59:387–415. doi: 10.1146/annurev.arplant.59.032607.092740

Flintham JE (2000) Different genetic components control coat-imposed and embryo-imposed dormancy in wheat. Seed Sci Res 10:43–50

Gerjets T, Scholefield D, Foulkes MJ, et al (2010) An analysis of dormancy, ABA responsiveness, after-ripening and pre-harvest sprouting in hexaploid wheat (*Triticum aestivum* L.) caryopses. J Exp Bot 61:597–607. doi: 10.1093/jxb/erp329

Gupta PK, Mir RR, Mohan A, Kumar J (2008) Wheat Genomics: Present Status and Future Prospects. Int J Plant Genomics. doi: 10.1155/2008/896451

Haley CS, Knott SA (1992) A simple regression method for mapping quantitative trait loci in line crosses using flanking markers. Heredity 69:315. doi: 10.1038/hdy.1992.131

Hamel L-P, Nicole M-C, Sritubtim S, et al (2006) Ancient signals: comparative genomics of plant MAPK and MAPKK gene families. Trends Plant Sci 11:192–198. doi: 10.1016/j.tplants.2006.02.007

Harrington SA, Cobo N, Karafiátová M, et al (2019) Identification of a dominant chlorosis phenotype through a forward screen of the *Triticum turgidum* cv. Kronos TILLING population. Front. Plant Sci., doi: 10.3389/fpls.2019.00963

Henry IM, Nagalakshmi U, Lieberman MC, et al (2014) Efficient Genome-Wide Detection and Cataloging of EMS-Induced Mutations Using Exome Capture and Next-Generation Sequencing. Plant Cell 26:1382–1397. doi: 10.1105/tpc.113.121590

Himi E, Maekawa M, Miura H, Noda K (2011) Development of PCR markers for *Tamyb10* related to *R-1*, red grain color gene in wheat. TAG Theor Appl Genet Theor Angew Genet 122:1561–1576. doi: 33-

Himi E, Mares DJ, Yanagisawa A, Noda K (2002) Effect of Grain Colour Gene (*R*) on Grain Dormancy and Sensitivity of the Embryo to Abscisic Acid (ABA) in Wheat. J Exp Bot 53:1569–1574. doi: 10.1093/jxb/erf005

IWGSC, Distelfeld A, Feuillet C, et al (2018) Shifting the limits in wheat research and breeding using a fully annotated reference genome. Science 361:eaar7191. doi: 10.1126/science.aar7191

IWGSC, Mayer KFX, Rogers J, et al (2014) A chromosome-based draft sequence of the hexaploid bread wheat (*Triticum aestivum*) genome. Science 345:1251788. doi: 10.1126/science.1251788

Jaiswal V, Mir RR, Mohan A, et al (2012) Association mapping for pre-harvest sprouting tolerance in common wheat (*Triticum aestivum* L.). Euphytica 188:89–102. doi: 10.1007/s10681-012-0713-1

Jiang M, Chu Z (2018) Comparative analysis of plant MKK gene family reveals novel expansion mechanism of the members and sheds new light on functional conservation. BMC Genomics 19:407. doi: 10.1186/s12864-018-4793-8

Joly-Lopez Z, Forczek E, Vello E, et al (2017) Abiotic Stress Phenotypes Are Associated with Conserved Genes Derived from Transposable Elements. Front Plant Sci 8:. doi: 10.3389/fpls.2017.02027

Kidwell KK, Demacon VL, Shelton GB, et al (2006a) Registration of ‘Otis’ Wheat. Crop Sci 46:1386–1387. doi: 10.2135/cropsci2005.06-0177

Kidwell KK, Shelton GB, Demacon VL, et al (2002) Registration of ‘Zak’ Wheat. Crop Sci 42:661–662. doi: 10.2135/cropsci2002.661a

Kidwell KK, Shelton GB, Demacon VL, et al (2006b) Registration of ‘Louise’ Wheat. Crop Sci 46:1384–1386. doi: 10.2135/cropsci2005.06-0176

Knetsch M, Wang M, Snaar-Jagalska BE, Heimovaara-Dijkstra S (1996) Abscisic Acid Induces Mitogen-Activated Protein Kinase Activation in Barley Aleurone Protoplasts. Plant Cell 8:1061–1067. doi: 10.1105/tpc.8.6.1061

Kosambi DD (1943) The Estimation of Map Distances from Recombination Values. Ann Eugen 12:172–175. doi: 10.1111/j.1469-1809.1943.tb02321.x

Krasileva KV, Vasquez-Gross HA, Howell T, et al (2017) Uncovering hidden variation in polyploid wheat. Proc Natl Acad Sci USA 114:E913–E921. doi: 10.1073/pnas.1619268114

Krzywinski M, Schein J, Birol İ, et al (2009) Circos: An information aesthetic for comparative genomics. Genome Res 19:1639–1645. doi: 10.1101/gr.092759.109

Kulwal P, Ishikawa G, Benscher D, et al (2012) Association mapping for pre-harvest sprouting resistance in white winter wheat. Theor Appl Genet 125:793–805. doi: 10.1007/s00122-012-1872-0

Langmead B, Trapnell C, Pop M, Salzberg SL (2009) Ultrafast and memory-efficient alignment of short DNA sequences to the human genome. Genome Biol 10:R25. doi: 10.1186/gb-2009-10-3-r25

Li H, Durbin R (2009) Fast and accurate short read alignment with Burrows–Wheeler transform. Bioinformatics 25:1754–1760. doi: 10.1093/bioinformatics/btp324

Li H, Handsaker B, Wysoker A, et al (2009a) The Sequence Alignment/Map format and SAMtools. Bioinforma Oxf Engl 25:2078–2079. doi: 10.1093/bioinformatics/btp352

Li R, Yu C, Li Y, et al (2009b) SOAP2: an improved ultrafast tool for short read alignment. Bioinformatics 25:1966–1967. doi: 10.1093/bioinformatics/btp336

Liu A, Gao F, Kanno Y, et al (2013a) Regulation of Wheat Seed Dormancy by After-Ripening Is Mediated by Specific Transcriptional Switches That Induce Changes in Seed Hormone Metabolism and Signaling. PLoS ONE 8:e56570. doi: 10.1371/journal.pone.0056570

Liu D, Li A, Mao X, Jing R (2014) Cloning and Characterization of *TaPP2AbB”-α*, a Member of the PP2A Regulatory Subunit in Wheat. PLoS ONE 9:e94430. doi: 10.1371/journal.pone.0094430

Liu S, Cai S, Graybosch R, et al (2008) Quantitative trait loci for resistance to pre-harvest sprouting in US hard white winter wheat Rio Blanco. Theor Appl Genet 117:691–699. doi: 10.1007/s00122-008-0810-7

Liu S, Sehgal SK, Li J, et al (2013b) Cloning and Characterization of a Critical Regulator for Preharvest Sprouting in Wheat. Genetics 195:263–273. doi: 10.1534/genetics.113.152330

Liu S, Yeh C-T, Tang HM, et al (2012) Gene Mapping via Bulked Segregant RNA-Seq (BSR-Seq). PLoS ONE 7:e36406. doi: 10.1371/journal.pone.0036406

Liu Y (2012) Roles of mitogen-activated protein kinase cascades in ABA signaling. Plant Cell Rep 31:1–12. doi: 10.1007/s00299-011-1130-y

Mares DJ, Mrva K (2014) Wheat grain preharvest sprouting and late maturity alpha-amylase. Planta 240:1167–1178. doi: 10.1007/s00425-014-2172-5

Martinez SA, Godoy J, Huang M, et al (2018) Genome-wide Association Mapping for Tolerance to Preharvest Sprouting and Low Falling Numbers in Wheat. Front Plant Sci 9:1–16. doi: 10.3389/fpls.2018.00141

Martinez SA, Schramm EC, Harris TJ, et al (2014) Registration of Zak Soft White Spring Wheat Germplasm with Enhanced Response to ABA and Increased Seed Dormancy. J Plant Regist 8:217–220. doi: 10.3198/jpr2013.09.0060crg

Martinez SA, Tuttle KM, Takebayashi Y, et al (2016) The wheat ABA hypersensitive *ERA8* mutant is associated with increased preharvest sprouting tolerance and altered hormone accumulation. Euphytica 212:229–245. doi: 10.1007/s10681-016-1763-6

McKibbin RS, Wilkinson MD, Bailey PC, et al (2002) Transcripts of *Vp-1* Homeologues Are Misspliced in Modern Wheat and Ancestral Species. Proc Natl Acad Sci USA 99:10203–10208. doi: 10.1073/pnas.152318599

Michelmore RW, Paran I, Kesseli RV (1991) Identification of markers linked to disease-resistance genes by bulked segregant analysis: a rapid method to detect markers in specific genomic regions by using segregating populations. Proc Natl Acad Sci USA 88:9828–9832. doi: 10.1073/pnas.88.21.9828

Mo Y, Howell T, Vasquez-Gross H, et al (2017) Mapping causal mutations by exome sequencing in a wheat TILLING population: a tall mutant case study. Mol Genet Genomics 1–15. doi: 10.1007/s00438-017-1401-6

Munkvold JD, Tanaka J, Benscher D, Sorrells ME (2009) Mapping quantitative trait loci for preharvest sprouting resistance in white wheat. Theor Appl Genet 119:1223–1235. doi: 10.1007/s00122-009-1123-1

Nakamura S, Abe F, Kawahigashi H, et al (2011) A Wheat Homolog of *MOTHER OF FT AND TFL1* Acts in the Regulation of Germination. Plant Cell 23:3215–3229. doi: 10.1105/tpc.111.088492

Nakamura S, Komatsuda T, Miura H (2007) Mapping diploid wheat homologues of *Arabidopsis* seed ABA signaling genes and QTLs for seed dormancy. TAG Theor Appl Genet Theor Angew Genet 114:1129–1139. doi: 10.1007/s00122-007-0502-8

Nakamura S, Pourkheirandish M, Morishige H, et al (2016) *Mitogen-Activated Protein Kinase Kinase 3* Regulates Seed Dormancy in Barley. Curr Biol 26:. doi: 10.1016/j.cub.2016.01.024

Nakamura S, Toyama T (2001) Isolation of a VP1 homologue from wheat and analysis of its expression in embryos of dormant and non☐ dormant cultivars. J Exp Bot 52:875–876. doi: 10.1093/jexbot/52.357.875

Nelson SK, Steber CM (2017) Transcriptional mechanisms associated with seed dormancy and dormancy loss in the gibberellin-insensitive sly1-2 mutant of Arabidopsis thaliana. PLOS ONE 12:e0179143. doi: 10.1371/journal.pone.0179143

Ogbonnaya FC, Imtiaz M, Ye G, et al (2008) Genetic and QTL analyses of seed dormancy and preharvest sprouting resistance in the wheat germplasm CN10955. Theor Appl Genet 116:891–902. doi: 10.1007/s00122-008-0712-8

Onate-Sánchez L, Vicente-Carbajosa J (2008) DNA-free RNA isolation protocols for Arabidopsis thaliana, including seeds and siliques. BMC Res Notes 1:93

Paterson AH, Sorrells ME, Obendorf RL (1989) Methods of Evaluation for Preharvest Sprouting Resistance in Wheat Breeding Programs. Can J Plant Sci 69:681–689. doi: 10.4141/cjps89-084

Pimentel H, Bray NL, Puente S, et al (2017) Differential analysis of RNA-seq incorporating quantification uncertainty. Nat Methods 14:687–690. doi: 10.1038/nmeth.4324

Poland JA, Brown PJ, Sorrells ME, Jannink J-L (2012) Development of High-Density Genetic Maps for Barley and Wheat Using a Novel Two-Enzyme Genotyping-by-Sequencing Approach. PLoS ONE 7:e32253. doi: 10.1371/journal.pone.0032253

Ramírez-González RH, Borrill P, Lang D, et al (2018) The transcriptional landscape of polyploid wheat. Science 361:eaar6089. doi: 10.1126/science.aar6089

Ramirez-Gonzalez RH, Segovia V, Bird N, et al (2015a) RNA-Seq bulked segregant analysis enables the identification of high-resolution genetic markers for breeding in hexaploid wheat. Plant Biotechnol J 13:613–624. doi: 10.1111/pbi.12281

Ramirez-Gonzalez RH, Uauy C, Caccamo M (2015b) PolyMarker: A fast polyploid primer design pipeline. Bioinformatics 31:2038–2039. doi: 10.1093/bioinformatics/btv069

Rodríguez MV, Barrero JM, Corbineau F, et al (2015) Dormancy in cereals (not too much, not so little): about the mechanisms behind this trait. Seed Sci Res 25:99–119. doi: 10.1017/S0960258515000021

Schramm EC (2010) Isolation of ABA-Response Mutants in Allohexaploid Bread Wheat (Triticum Aestivum L.): Drawing Connections to Grain Dormancy, Preharvest Sprouting, and Drought Tolerance. Ph.D., Washington State University

Schramm EC, Abellera JC, Strader LC, et al (2010) Isolation of ABA-responsive mutants in allohexaploid bread wheat (*Triticum aestivum* L.): Drawing connections to grain dormancy, preharvest sprouting, and drought tolerance. Plant Sci 179:620–629. doi: 10.1016/j.plantsci.2010.06.004

Schramm EC, Nelson SK, Kidwell KK, Steber CM (2013) Increased ABA sensitivity results in higher seed dormancy in soft white spring wheat cultivar ‘Zak.’ Theor Appl Genet 126:791–803. doi: 10.1007/s00122-012-2018-0

Schramm EC, Nelson SK, Steber CM (2012) Wheat ABA-insensitive mutants result in reduced grain dormancy. Euphytica 188:35–49

Shewry PR (2009) Wheat. J Exp Bot 60:1537–1553. doi: 10.1093/jxb/erp058

Shibata W, Banno H, Ito Y, et al (1995) A tobacco protein kinase, NPK2, has a domain homologous to a domain found in activators of mitogen-activated protein kinases (MAPKKs). Mol Gen Genet MGG 246:401–410. doi: 10.1007/BF00290443

Shorinola O, Balcárková B, Hyles J, et al (2017) Haplotype Analysis of the Pre-harvest Sprouting Resistance Locus Phs-A1 Reveals a Causal Role of TaMKK3-A in Global Germplasm. Front Plant Sci 8:1–14. doi: 10.3389/fpls.2017.01555

Shorinola O, Bird N, Simmonds J, et al (2016) The wheat *Phs-A1* pre-harvest sprouting resistance locus delays the rate of seed dormancy loss and maps 0.3 cM distal to the *PM19* genes in UK germplasm. J Exp Bot 67:4169–4178. doi: 10.1093/jxb/erw194

Smith SM, Maughan PJ (2015) SNP Genotyping Using KASPar Assays. In: Plant Genotyping. Humana Press, New York, NY, pp 243–256

Stekhoven DJ, Bühlmann P (2012) MissForest--non-parametric missing value imputation for mixed-type data. Bioinforma Oxf Engl 28:112–118. doi: 10.1093/bioinformatics/btr597

Takahashi F, Yoshida R, Ichimura K, et al (2007) The Mitogen-Activated Protein Kinase Cascade MKK3–MPK6 Is an Important Part of the Jasmonate Signal Transduction Pathway in Arabidopsis. Plant Cell 19:805–818. doi: 10.1105/tpc.106.046581

Thole JM, Strader LC (2015) Next-generation sequencing as a tool to quickly identify causative EMS-generated mutations. Plant Signal Behav 10:1–4. doi: 10.1080/15592324.2014.1000167

Torada A, Amano Y (2002) Effect of seed coat color on seed dormancy in different environments. Euphytica 126:99–105. doi: 10.1023/A:1019603201883

Torada A, Koike M, Ogawa T, et al (2016) A Causal Gene for Seed Dormancy on Wheat Chromosome 4A Encodes a MAP Kinase Kinase. Curr Biol 26:782–787. doi: 10.1016/j.cub.2016.01.063

Trick M, Adamski NM, Mugford SG, et al (2012) Combining SNP discovery from next-generation sequencing data with bulked segregant analysis (BSA) to fine-map genes in polyploid wheat. BMC Plant Biol 12:14. doi: 10.1186/1471-2229-12-14

Truong HT, Ramos AM, Yalcin F, et al (2012) Sequence-Based Genotyping for Marker Discovery and Co-Dominant Scoring in Germplasm and Populations. PLOS ONE 7:e37565. doi: 10.1371/journal.pone.0037565

Tuttle KM, Martinez SA, Schramm EC, et al (2015) Grain dormancy loss is associated with changes in ABA and GA sensitivity and hormone accumulation in bread wheat, *Triticum aestivum* (L.). Seed Sci Res 25:179–193. doi: 10.1017/S0960258515000057

Uauy C, Wulff BBH, Dubcovsky J (2017) Combining Traditional Mutagenesis with New High-Throughput Sequencing and Genome Editing to Reveal Hidden Variation in Polyploid Wheat. Annu Rev Genet 51:435–454. doi: 10.1146/annurev-genet-120116-024533

Van Ooijen JW (2006) JoinMap (R) 4, Software for the calculation of genetic linkage maps in experimental populations. B.V. Kyazma, Wageningen, Netherlands

Walker-Simmons MK (1987) ABA Levels and Sensitivity in Developing Wheat Embryos of Sprouting Resistant and Susceptible Cultivars. Plant Physiol 84:61–66. doi: 10.1104/pp.84.1.61

Wickham H (2009) ggplot2: Elegant Graphics for Data Analysis. Springer-Verlag New York

Xiong L, Yang Y (2003) Disease Resistance and Abiotic Stress Tolerance in Rice Are Inversely Modulated by an Abscisic Acid–Inducible Mitogen-Activated Protein Kinase. Plant Cell 15:745–759. doi: 10.1105/tpc.008714

Xu Q, Truong TT, Barrero JM, et al (2016) A role for jasmonates in the release of dormancy by cold stratification in wheat. J Exp Bot erw172. doi: 10.1093/jxb/erw172

Yan L, Loukoianov A, Tranquilli G, et al (2003) Positional cloning of the wheat vernalization gene VRN1. Proc Natl Acad Sci USA 100:6263–6268. doi: 10.1073/pnas.0937399100

Yu G, Hatta A, Periyannan S, et al (2017) Isolation of Wheat Genomic DNA for Gene Mapping and Cloning. In: Wheat Rust Diseases. Humana Press, New York, NY, pp 207–213

Zanetti S, Winzeler M, Keller M, et al (2000) Genetic analysis of pre-harvest sprouting resistance in a wheat × spelt cross. Crop Sci 40:1406–1417. doi: 10.2135/cropsci2000.4051406x

Zhou Y, Tang H, Cheng M-P, et al (2017) Genome-Wide Association Study for Pre-harvest Sprouting Resistance in a Large Germplasm Collection of Chinese Wheat Landraces. Front Plant Sci 8:. doi: 10.3389/fpls.2017.00401

Zuo J, Lin C-T, Cao H, et al (2019) Genome-wide association study and quantitative trait loci mapping of seed dormancy in common wheat (*Triticum aestivum* L.). Planta 250:187–198. doi: 10.1007/s00425-019-03164-9

