## Supplemental Material 1 for "Exome sequencing of bulked segregants identified a novel *TaMKK3-A* allele linked to the wheat *ERA8* ABA-hypersensitive germination phenotype"

Fig. S1 - Crossing strategy to generate the *ERA8* backcross mapping population
Fig. S2 - WT and *ERA8* parental seed germination assay
Fig. S3 - Insertions or deletions between WT and *ERA8*
Fig. S4 - ABA response to number of QTL in the Louise/Zak*ERA8* RILs
Fig. S5 - QTL analysis of Louise/Zak*ERA8* heading date and height
Fig. S6 - WT and *ERA8* coding sequence of *TaMKK3-A*

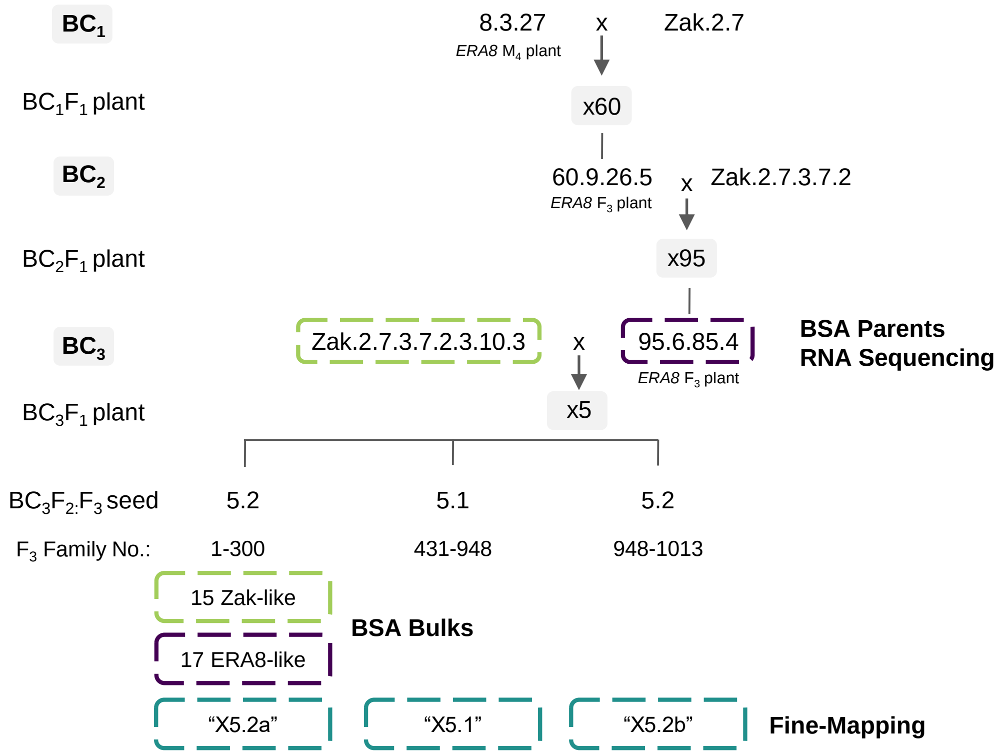


**Fig. S1** Crossing strategy to generate the *ERA8* backcross mapping population. Zak WT parent (Zak2.7.3.7.2.3.10.3), the *ERA8* parent (95.6.85.4), *ERA8*-like bulk, and Zak-like bulk samples indicated by the dashed **green** and **purple** boxes were used in the bulked segregant analysis (BSA) exome sequencing. The $BC_3F_2$:$F_3$ lines used in the fine mapping study are indicated by the dashed **blue** boxes. The parent on the left of the ‘x’ is the female plant used in the cross.

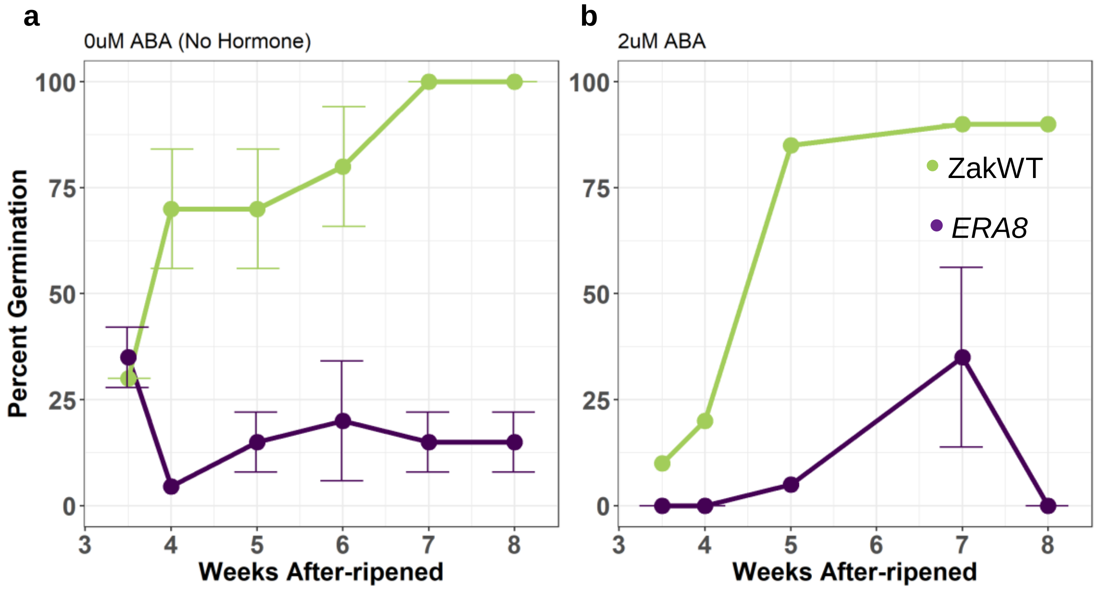


**Fig. S2** WT and ERA8 parental seed germination assay. An after-ripening time course of the *ERA8* (purple) and ZakWT (green) parents from the Zak/Zak*ERA8* was conducted without ABA (No Hormone; **a**) and with 2 μM ABA (**b**). Percent germination after 5 days of imbibition at 30° was calculated from germination assays of 20 whole seeds each. The five week after-ripened time point was chosen for bulk segregant analysis.

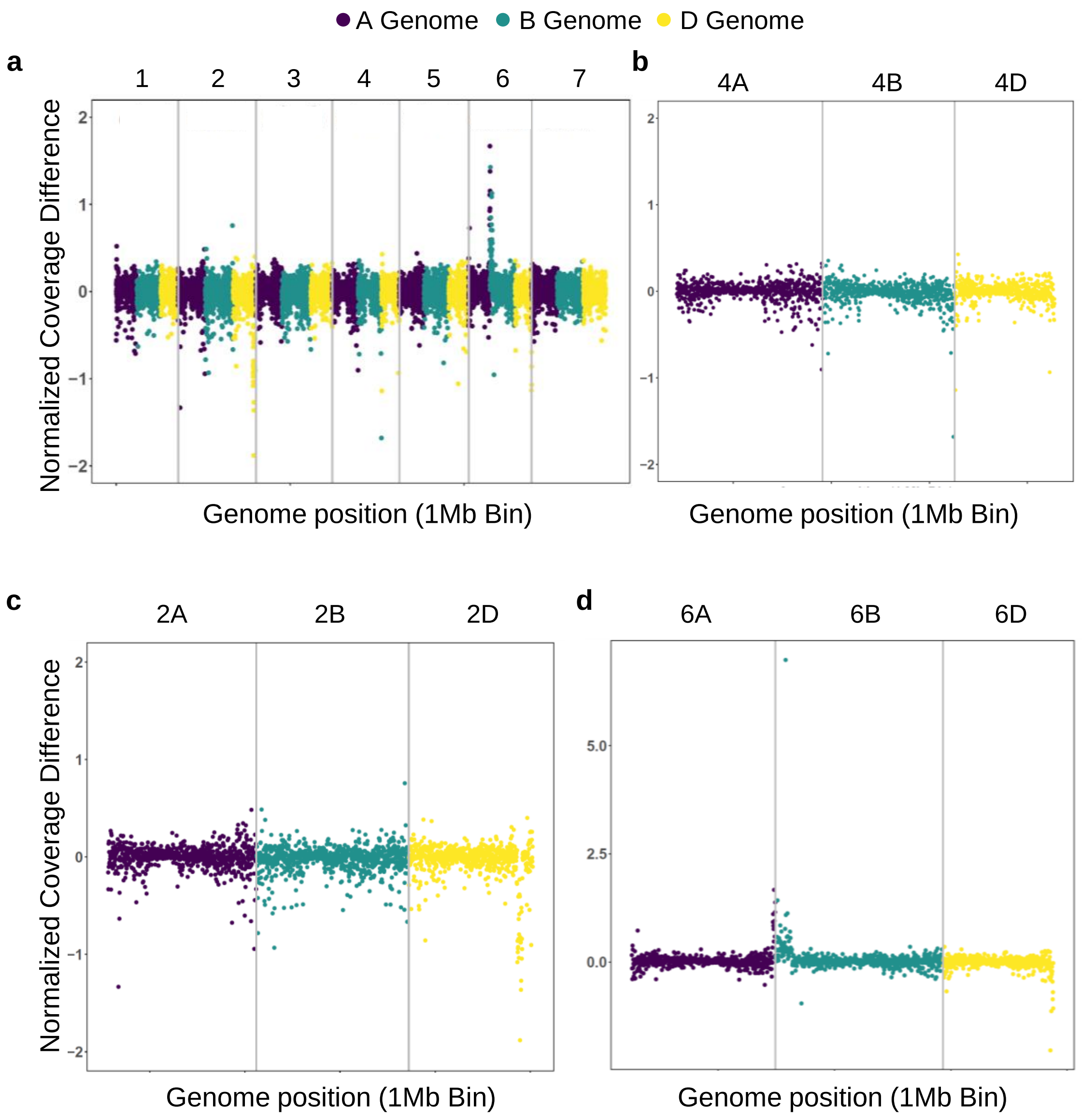


**Fig. S3** Insertions or deletions between WT and *ERA8*. To identify potential insertions and deletions, read coverages in Zak WT were compared to *ERA8* across the: **a**) whole genome; **b**) group 4 chromosomes; **c**) group 2 chromosomes, and **d**) group 6 chromosomes. Insertions and deletions in *ERA8* are indicated by a positive or negative difference, respectively. The A genome is shown in purple, the B genome in green, and the D genome in yellow.

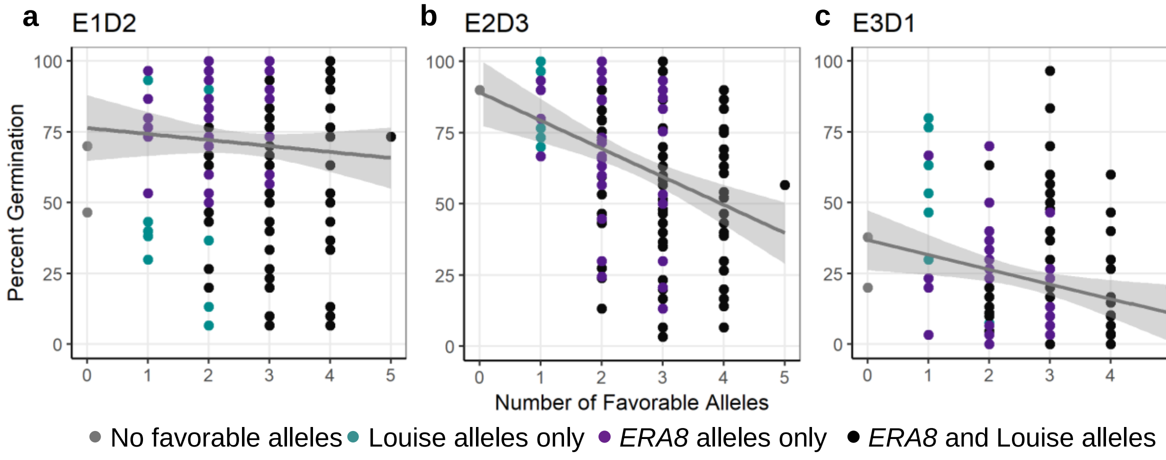


**Fig. S4** ABA response to number of QTL in the Louise/Zak*ERA8* RILs. The number of favorable alleles that increase ABA sensitivity is compared against percent germination. Environments E1, E2, and E3 are shown after **a**) two, **b**) three, and **c**) one day of imbibition, respectively. Individual RILs are shown to have no tolerant alleles (grey), only the Louise (blue) or *ERA8* (purple) parent contributes to the tolerant allele, or both parents (black) contribute to the tolerant alleles. A simple linear regression line is shown using `geom_smooth(method = lm)` in the ggplot R package.

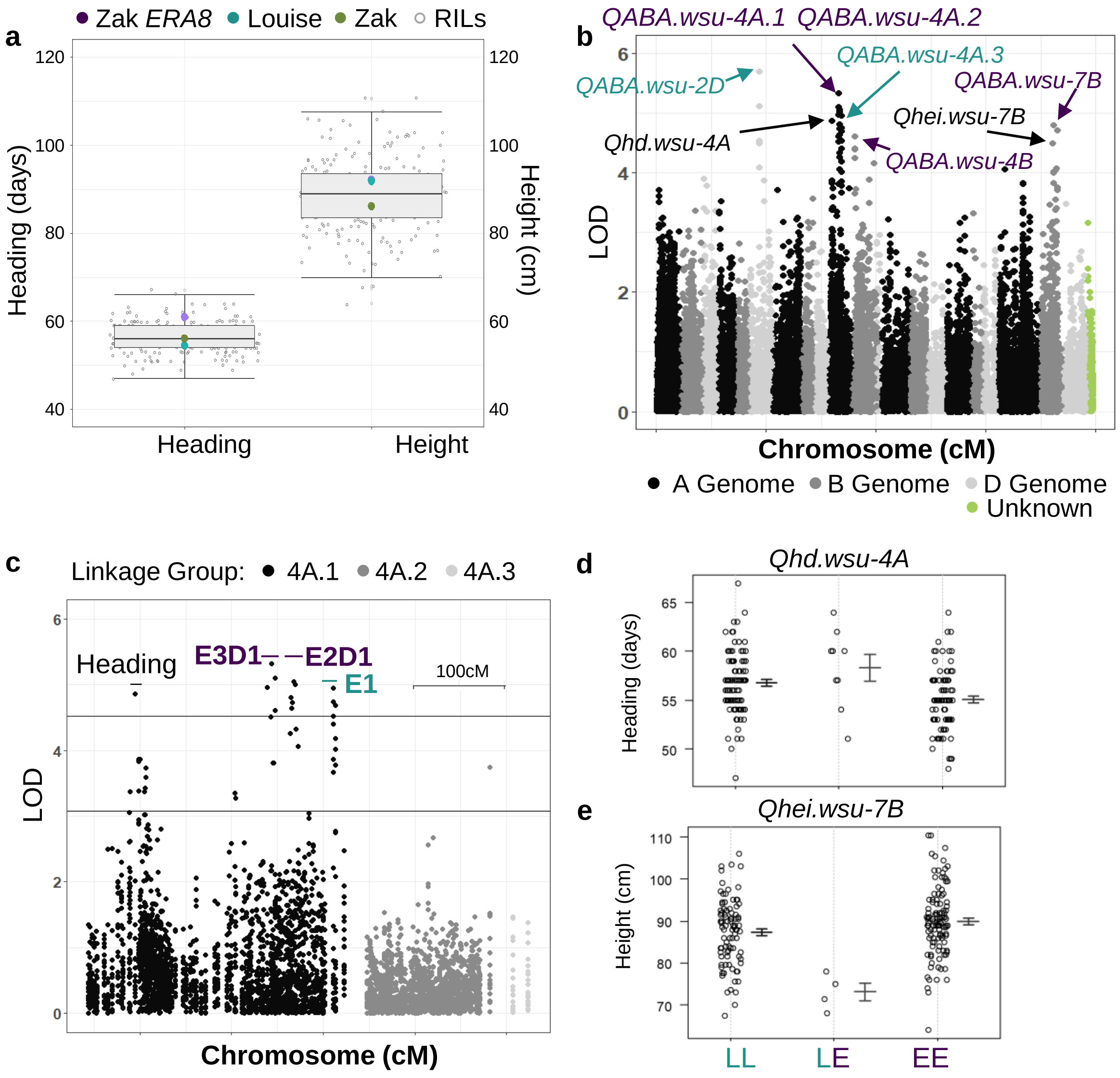


**Fig. S5** – QTL analysis of heading and height data. LOD scores for **a**) heading and height of the Louise/Zak*ERA8* RIL population are shown across **b**) all chromosomes (cM) and **c**) only for chromosome 4A.  are shown. Significant QTL for height (*Qhei.wsu;* black), days to heading (*Qhd.wsu;* black), and ABA sensitivity (*QABA.wsu*), with a threshold of $p <$ 0.10 are shown. QTL where *ERA8* or Louise contributes to the ABA sensitivity are shown as purple or blue, respectively. Alleles were compared between heading date (**d**; days) and height (**e;** cm) differences in the RIL population separated by Louise (LL) and *ERA8* (EE) alleles. Note that height was taken from a single plant and not multiple plants across a plot.

**a** TaMKK3-A C660A mutation   EMS G1093A mutation

**Zak WT**

ATGTCGGGGCTAGAGGAGTTGAAGAAGAAGCTGCAGCCCTTGCTGTTCGACGACACGGACAAGGGCGGCGTCAGTACCCGGGTTCCCTTCCCGGAGGATACATGCGATTCCTATGT
GGTGTCTGATGGTGGAACGATAAATTTACTGAGTAGATCGTTTGGTGAGTATAACATCAATGAGCATGGCTTTCATAAGCGAAGCACCGGACCAGAAGAGCCAGATAGCGGTGAGA
AGGCATACCGATGTGCCTCTGAAGACATGCACATATTTGGTCCCATTGGGAATGGAGCAAGCAGTGTCGTGCAGAGAGCTATTTTCATACCAGTTCATCGAATTCTGGCCTTGAAG
AAGATAAACATATTTGAGAAGGAGAAAAGGCAACAAATTCTTAATGAGATGAGAACATTATGTGAAGCAAGTTGCTATCCTGGTTTAGTTGAATTCCAGGGTGCATTTTACATGCC
TGATTCTGGACAAATAAGCATTGCCCTTGAATACATGGATGGTGGCTCTTTAGCAGACGTTATAAAAGTCAAGAAATCAATACCAGAGCCAGTTCTTGCACATATGCTACAGAAAG
TGTTACTTGGCCTGAAGTACTTGCATGAAGTAAGACATCTAGTGCATAGAGATCTAAAGCCAGCAAATTTACTGGTAAAACTTAAGGGCGAAGCAAAAATTACAGATTTTGGTGTA
AGTGCTGGTTTGGACAATACAATGGCTATGTGTGCTACCTTTGTAGGCACTGTGACATATATGTCACCTGAGAGAATCCGCAATGAGAACTACTCCTATGCTGCTGATATTTGGAG
TCTTGGACTAACGATATTGGAGTGTGCTACTGGTAAATTTCCATATAATGTCAACGAAGGCCCAGCCAATCTCATGCTACAGATACTCGATGATCCATCACCAACACCACCAGAAG
ATGCCTATACACCAGAATTTTGTTCCTTCATAAATGATTGCTTGAGGAAAGATGCTGATGCAAGGCCTACATGCGAGCAGCTTTTGTCACACGCATTTATCAAGAGGTATGAGCAA
ACTGGTGTGGACTTGGCAGCATATGTCAGGGGTGTTGTTAATCCAACAGAGAGATTAAAGCAAATAGCAGAGATGCTTGCTGTTCATTATTACCTCCTTTTTAATGGCTCTGAAGG
ACCTTGGAATTATATGAAGACATTCTACAAGGAGGAATCATCTTTCAGTTTTTCAGGGAATGTGTATGTCGGACAAAGTGCCATATTTGATACTTTATCAAATATAAGAAAGAAGT
TGAAAGGTGACCGGCCTAGAGAAAAAATTGTTCATGTTGTTGAGAAGCTACATTGCCGCGCGAATGGGGAAACAGAAATTGCTATCCGTGTGTCTGGATCATTCATCACGGGCAAC
CAATTCCTGATATTTGGTGAAGGGTTGCAAGCTGAAGGGATGCCCAGCTTGGAGGAAATTGATATTGACATTCCAAGCAAGCGGGTTGGCCAGTTCCGGGAGCAGTTTACCGTGCA
TCCAGGGACTTCCATGGGATGCTACTACATAGCAAAGCAAGATCTCTACATCGTCCAGTCATGA

***ERA8***

ATGTCGGGGCTAGAGGAGTTGAAGAAGAAGCTGCAGCCCTTGCTGTTCGACGACACGGACAAGGGCGGCGTCAGTACCCGGGTTCCCTTCCCGGAGGATACATGCGATTCCTATGT
GGTGTCTGATGGTGGAACGATAAATTTACTGAGTAGATCGTTTGGTGAGTATAACATCAATGAGCATGGCTTTCATAAGCGAAGCACCGGACCAGAAGAGCCAGATAGCGGTGAGA
AGGCATACCGATGTGCCTCTGAAGACATGCACATATTTGGTCCCATTGGGAATGGAGCAAGCAGTGTCGTGCAGAGAGCTATTTTCATACCAGTTCATCGAATTCTGGCCTTGAAG
AAGATAAACATATTTGAGAAGGAGAAAAGGCAACAAATTCTTAATGAGATGAGAACATTATGTGAAGCAAGTTGCTATCCTGGTTTAGTTGAATTCCAGGGTGCATTTTACATGCC
TGATTCTGGACAAATAAGCATTGCCCTTGAATACATGGATGGTGGCTCTTTAGCAGACGTTATAAAAGTCAAGAAATCAATACCAGAGCCAGTTCTTGCACATATGCTACAGAAAG
TGTTACTTGGCCTGAAGTACTTGCATGAAGTAAGACATCTAGTGCATAGAGATCTAAAGCCAGCAAATTTACTGGTAAAACTTAAGGGCGAAGCAAAAATTACAGATTTTGGTGTA
AGTGCTGGTTTGGACAATACAATGGCTATGTGTGCTACCTTTGTAGGCACTGTGACATATATGTCACCTGAGAGAATCCGCAATGAGAACTACTCCTATGCTGCTGATATTTGGAG
TCTTGGACTAACGATATTGGAGTGTGCTACTGGTAAATTTCCATATAATGTCAACGAAGGCCCAGCCAATCTCATGCTACAGATACTCGATGATCCATCACCAACACCACCAGAAG
ATGCCTATACACCAGAATTTTGTTCCTTCATAAATGATTGCTTGAGGAAAGATGCTGATGCAAGGCCTACATGCGAGCAGCTTTTGTCACACGCATTTATCAAGAGGTATGAGCAA
ACTGGTGTGGACTTGGCAGCATATGTCAGGGGTGTTGTTAATCCAACAAAGAGATTAAAGCAAATAGCAGAGATGCTTGCTGTTCATTATTACCTCCTTTTTAATGGCTCTGAAGG
ACCTTGGAATTATATGAAGACATTCTACAAGGAGGAATCATCTTTCAGTTTTTCAGGGAATGTGTATGTCGGACAAAGTGCCATATTTGATACTTTATCAAATATAAGAAAGAAGT
TGAAAGGTGACCGGCCTAGAGAAAAAATTGTTCATGTTGTTGAGAAGCTACATTGCCGCGCGAATGGGGAAACAGAAATTGCTATCCGTGTGTCTGGATCATTCATCACGGGCAAC
CAATTCCTGATATTTGGTGAAGGGTTGCAAGCTGAAGGGATGCCCAGCTTGGAGGAAATTGATATTGACATTCCAAGCAAGCGGGTTGGCCAGTTCCGGGAGCAGTTTACCGTGCA
TCCAGGGACTTCCATGGGATGCTACTACATAGCAAAGCAAGATCTCTACATCGTCCAGTCATGA

**b** Catalytic domain of Plant dual-specificity Mitogen-Activated Protein Kinase Kinases   Nuclear transport factor 2 (NTF2)   EMS mutation

**Zak WT**

MSGLEELKKKLQPLLFDDTDKGGVSTRVPFPEDTCDSYVVSDGGTINLLSRSFGEYNINEHGFHKRSTGPEEPDSGEKAYRCASEDMHIFGPIGNGASSVVQRAIFIPVHRILALK
KINIFEKEKRQQILNEMRTLCEASCYPGLVEFQGAFYMPDSGQISIALEYMDGGSLADVIKVKKSIPEPVLAHMLQKVLLGLKYLHEVRHLVHRDLKPANLLVKLKGEAKITDFGV
SAGLDNTMAMCATFVGTVTYMSPERIRNENYSYAADIWSLGLTILECATGKFPYNVNEGPANLMLQILDDPSPTPPEDAYTPEFCSFINDCLRKDADARPTCEQLLSHAFIKRYEQ
TGVDLAAYVRGVVNPTERLKQIAEMLAVHYYLLFNGSEGPWNYMKTFYKEESSFSFSGNVYVGQSAIFDTLSNIRKKLKGDRPREKIVHVVEKLHCRANGETEIAIRVSGSFITGN
QFLIFGEGLQAEGMPSLEEIDIDIPSKRVGQFREQFTVHPGTSMGCYYIAKQDLYIVQS-

***ERA8***

MSGLEELKKKLQPLLFDDTDKGGVSTRVPFPEDTCDSYVVSDGGTINLLSRSFGEYNINEHGFHKRSTGPEEPDSGEKAYRCASEDMHIFGPIGNGASSVVQRAIFIPVHRILALK
KINIFEKEKRQQILNEMRTLCEASCYPGLVEFQGAFYMPDSGQISIALEYMDGGSLADVIKVKKSIPEPVLAHMLQKVLLGLKYLHEVRHLVHRDLKPANLLVKLKGEAKITDFGV
SAGLDNTMAMCATFVGTVTYMSPERIRNENYSYAADIWSLGLTILECATGKFPYNVNEGPANLMLQILDDPSPTPPEDAYTPEFCSFINDCLRKDADARPTCEQLLSHAFIKRYEQ
TGVDLAAYVRGVVNPTKRLKQIAEMLAVHYYLLFNGSEGPWNYMKTFYKEESSFSFSGNVYVGQSAIFDTLSNIRKKLKGDRPREKIVHVVEKLHCRANGETEIAIRVSGSFITGN
QFLIFGEGLQAEGMPSLEEIDIDIPSKRVGQFREQFTVHPGTSMGCYYIAKQDLYIVQS-

**Fig. S6** –WT and *ERA8* coding sequence of *TaMKK3-A*. **a**) Coding region nucleotide sequences include the *TaMKK3-A* C660A mutation (blue) found in Torada et al. (2016) and the *TaMKK3-A* G1093A EMS mutation (purple) found between Zak WT and *ERA8*. **b**) The protein sequence of *TaMKK3-A* contains a Catalytic domain of Plant dual-specificity Mitogen-Activated Protein Kinase Kinases (light blue) and a Nuclear transport factor 2 (NTF2; green). Note this is reverse complement relative to the reference scaffold.
