## Supplemental Material 2 for "Exome sequencing of bulked segregants identified a novel *TaMKK3-A* allele linked to the wheat *ERA8* ABA-hypersensitive germination phenotype"

Table S1 - Summary of Zak/Zak*ERA8* ABA phenotyping conditions
Table S2 - Chi-squared analysis of Zak/Zak*ERA8* BC3F2:3 X5.2
Table S3 - Exome capture sequencing quality and statistics
Table S4 - Primers created from the GBS and exome capture identified SNPs
Table S5 - Chi-squared analysis of Zak/Zak*ERA8* BC3F2:3 X5.1 and X5.2
Table S6 - Louise/Zak*ERA8* linkage group summary
Table S7 - Genes in the *ERA8* 4A region
Table S8 - Summary of Louise/Zak*ERA8* ABA phenotyping conditions
Table S9 - Significant QTL of ABA sensitivity, height, and heading date for Louise/Zak*ERA8*
Table S10 - Genes differentially expressed between Zak and *ERA8*

**Table S1** ABA sensitivity for the Zak/Zak*ERA8* $BC_3F_{2:3}$ population across different seed lots

| Location | Year | n [a] | b rep [b] | ABA (μM) | AR (days) | Imb. [c] (days) | Mean ± SD [d] PG | GI | Storage in -20 °C (months) |
|---|---|---|---|---|---|---|---|---|---|
| 5.2 BSA | | 15 | 3 | 2 | 35 | 5 | 12.6 ± 8.5 | - | 4 |
| | | 17 | 3 | 2 | 35 | 5 | 79.9 ± 13 | - | 4 |
| | | *ZakERA8* | 16 | 2 | 35 | 5 | 8.3 ± 7.2 | - | 4 |
| | | *Zak* | 16 | 2 | 35 | 5 | 90.8 ± 15 | - | 4 |
| 5.2a | | 242 | 2 | 2 | 35 | 5 | 42.6 ± 21 | 0.21 ± 0.1 | 17 |
| | | *ZakERA8* | 8 | 2 | 35 | 5 | 8.3 ± 4.3 | 0.03 ± 0.0 | 17 |
| | | *Zak* | 8 | 2 | 35 | 5 | 90.8 ± 12 | 0.55 ± 0.0 | 17 |
| 5.1 | | 122 | 1 | 10 | 42, 59 | 4 | 43.2 ± 25 | 0.29 ± 0.1 | 6 |
| | | *ZakERA8* | 5 | 10 | 42, 59 | 4 | 54.3 ± 15 | 0.14 ± 0.1 | 6 |
| | | *Zak* | 5 | 10 | 42, 59 | 4 | 89.3 ± 5.5 | 0.40 ± 0.1 | 6 |
| 5.2b | | 60 | 3 | 5 | 41 | 4 | 68.5 ± 17 | 0.42 ± 0.1 | 3 |
| | | *ZakERA8* | 12 | 5 | 41 | 4 | 12.7 ± 5.2 | 0.08 ± 0.0 | 3 |
| | | *Zak* | 12 | 5 | 41 | 4 | 90.5 ± 3.5 | 0.62 ± 0.0 | 3 |

[a] number (n) of backcross lines tested

[b] technical replicates per backcross or parental line.

[c] The number of days imbibed that was used for bulk segregant analysis (BSA) or fine mapping

[d] Raw mean and standard deviation (SD) across the
backcross population, Zak, or *ERA8*.

**Table S2** Segregation analysis of $BC_3F_2$ seed germination on 2uM ABA

| Genotype | n[a] | Gen.[b] | Germ.[c] | Not Germ. [c] | $\chi^2$ | | | p-value | | |
|---|---|---|---|---|---|---|---|---|---|---|
| | | | | | **3:1** | **1:3** | **1:2:1** | **3:1** | **1:3** | **1:2:1** |
| 5.2a segregating | 300 | $BC_3F_2$ | 154 | 146 | 140.7 | 36.6 | 1.62 | <0.001 | <0.001 | **0.20** |
| +/+ | 60 | parent[e] | 54 | 6 | | | | | | |
| *ERA8/ERA8* | 60 | parent[e] | 0 | 60 | | | | | | |
| +/*ERA8* | 8 | $BC_2F_1$ | 4 | 4 | | | | | | |
| $F_2$ Expected[d] | 300 | | | | 68 | 203 | 143 | | | |

[a] Number of seeds tested for germination, after-ripened for 5 weeks past physiological maturity; df = 1

[b] Generation of seeds tested

[c] Number of seeds that had germinated (Germ.) and not germinated (Not Germ.) after 5 days of imbibition on 2μM (+/-) ABA.

[d] Number of seeds expected to germinate after 5 days of imbibition for each single gene segregation ratio.

[e] Zak (+/+) and *ERA8* (*ERA8*/*ERA8*) parental lines were used to generate cross 5; Parents were grown at the same time as the $F_1$ plants.

**Table S3** Sequencing read count and quality statistics from exome capture

| Sample Name | Number of Reads | Mean Q30 to base Read 1 | Mean Q30 to base Read 2 |
|---|---|---|---|
| *ERA8* Parent | 60,514,766 | 125 | 125 |
| *ERA8*_Bulk | 57,332,604 | 126 | 125 |
| WT Parent | 48,792,615 | 126 | 125 |
| WT_Bulk | 67,097,197 | 126 | 125 |

**Table S4** *ERA8* primers created from the GBS and Exome capture identified SNPs

| Name | Allele | Sequence | Tag | Source |
|---|---|---|---|---|
| SNP_1 | WT | GAAGGTGACCAAGTTCATGCTaCtctttctcatgagctctttgaG | FAM | Exome |
| | *ERA8* | GAAGGTCGGAGTCAACGGATTaCtctttctcatgagctctttgaA | HEX | |
| | C | tcgcgttcatcatctacctG | - | |
| SNP_2 | WT | GAAGGTGACCAAGTTCATGCTctgaaatgCctgcaagatgaG | FAM | Exome |
| | *ERA8* | GAAGGTCGGAGTCAACGGATTctgaaatgCctgcaagatgaA | HEX | |
| | C | ggaaggccaaaatagcgacttA | - | |
| SNP_4 | WT | GAAGGTGACCAAGTTCATGCTaaactcgacGaaaTtgcCaTC | FAM | Exome |
| | *ERA8* | GAAGGTCGGAGTCAACGGATTaaactcgacGaaaTtgcCaTT | HEX | |
| | C | aatcaacaCcattttcatttcacaG | - | |
| SNP_5 | WT | GAAGGTGACCAAGTTCATGCTgtggagtgTctgcaagatcG | FAM | Exome |
| | *ERA8* | GAAGGTCGGAGTCAACGGATTgtggagtgTctgcaagatcA | HEX | |
| | C | tatgcactcaccacgccA | - | |
| SNP_6 | WT | GAAGGTGACCAAGTTCATGCTCacaaactgtgctatcccgG | FAM | Exome |
| | *ERA8* | GAAGGTCGGAGTCAACGGATTCacaaactgtgctatcccgA | HEX | |
| | C | tggtagcagttgtcttgacC | - | |
| SNP_7 | WT | GAAGGTGACCAAGTTCATGCTggcttgcgggtaaaagagG | FAM | Exome |
| | *ERA8* | GAAGGTCGGAGTCAACGGATTggcttgcgggtaaaagagA | HEX | |
| | C | tctttgttttataattgcagtgcaC | - | |
| SNP_8 | WT | GAAGGTGACCAAGTTCATGCTtgTtAGatgtGaccttacctctcaG | FAM | Exome |
| | *ERA8* | GAAGGTCGGAGTCAACGGATTtgTtAGatgtGaccttacctctcaA | HEX | |
| | C | cgagatgagcaaggaggtgg | - | |
| SNP_9 | WT | GAAGGTGACCAAGTTCATGCTCggttctgcgttaccccatG | FAM | Exome |
| | *ERA8* | GAAGGTCGGAGTCAACGGATTCggttctgcgttaccccatA | HEX | |
| | C | agcaaatgCcgCggatcG | - | |
| SNP_10 | WT | GAAGGTGACCAAGTTCATGCTCggatcacgaTcgccttctcC | FAM | Exome |
| | *ERA8* | GAAGGTCGGAGTCAACGGATTCggatcacgaTcgccttctcT | HEX | |
| | C | TgTgctcctcGtcCtGAC | - | |
| SNP_17 | WT | GAAGGTGACCAAGTTCATGCTCCTCTGCTATTTGCTTTAATCTCTc | FAM | Exome |
| | *ERA8* | GAAGGTCGGAGTCAACGGATTCCTCTGCTATTTGCTTTAATCTCTt | VIC | |
| | C | GGACTTGGCAGCATATGTCA | - | |
| SNP_20 | WT | GAAGGTGACCAAGTTCATGCTCtcctgctcctgcttccgG | FAM | Exome |
| | *ERA8* | GAAGGTCGGAGTCAACGGATTCtcctgctcctgcttccgA | HEX | |
| | C | cggcctcacttgcagaaaac | - | |
| SNP_29 | WT | GAAGGTGACCAAGTTCATGCTgtgtacgcGcgCtactgC | FAM | Exome |
| | *ERA8* | GAAGGTCGGAGTCAACGGATTgtgtacgcGcgCtactgT | HEX | |
| | C | ccatgatctccagcgacagA | - | |
| SNP_30 | WT | GAAGGTGACCAAGTTCATGCTggtataaacatcagaatccctgtcC | FAM | Exome |
| | *ERA8* | GAAGGTCGGAGTCAACGGATTggtataaacatcagaatccctgtcT | HEX | |
| | C | ccggcatcttcTGtattaacatacA | - | |
| PHS1[a] | Res | GAAGGTGACCAAGTTCATGCTTTTTGCTTCGCCCTTAAGG | FAM | - |
| | Susc | GAAGGTCGGAGTCAACGGATTTTTTTGCTTCGCCCTTAAGT | HEX | |
| | C | GCATAGAGATCTAAAGCCAGCA | - | |
| A7575 | WT | GAAGGTCGGAGTCAACGGATTCaAActacacactcgtcgggA | HEX | GBS |
| | *ERA8* | GAAGGTGACCAAGTTCATGCTCaAActacacactcgtcgggG | FAM | |
| | C | ccctgcagcagaggacatC | - | |
| A7946 | *ERA8* | GAAGGTCGGAGTCAACGGATTCaAactacacactcgtcgggA | HEX | GBS |
| | Ref | GAAGGTGACCAAGTTCATGCTCaAactacacactcgtcgggG | FAM | |
| | C | ccctgcagcagaggacatC | - | |

[a] PHS1 KASP primers were developed by Shorinola et al. (2017)

[b] KASP tags (FAM and HEX) are already included in the primer sequence and indicated by grey text

**Table S5** Segregation analysis of Zak/*ZakERA8* $BC_3F_2$ X5.1, X5.2, and X5.3 germination on ABA

| **Genotype** | **n**[a] | **Gen.**[b] | **Germ.**[c] | **Not Germ.**[c] | **$\chi^2$** | | | **p-value** | | |
|---|---|---|---|---|---|---|---|---|---|---|
| | | | | **$F_2$** | **3:1** | **1:3** | **1:2:1** | **3:1** | **1:3** | **1:2:1** |
| | | | | **$F_3$** | **0.625:0.375** | **0.375:0.625** | **0.375:0.25:0.375** | **0.625:0.375** | **0.375:0.625** | **0.375:0.25:0.375** |
| 56d AR \| 10uM ABA \| 4d imb. | | | | | | | | | | |
| **X5.1** | 1311 | $BC_3F_3$ | 588 | 723 | 18.6 | 39.8 | 0.08 | <0.001 | <0.001 | 0.78 |
| WT/WT | 54 | parent[e] | 39 | 15 | | | | | | |
| *ERA8/ERA8* | 60 | parent[e] | 9 | 51 | | | | | | |
| WT/*ERA8* | 8 | $BC_2F_1$ | 4 | 4 | | | | | | |
| $F_3$ Expected[d] | 1311 | | | | 666 | 478 | 593 | | | |
| 35d AR \| 2uM ABA \| 4d imb. | | | | | | | | | | |
| **X5.2a** | 7361 | $BC_3F_3$ | 3974 | 3387 | 107.0 | 677.3 | 55.3 | <0.001 | <0.001 | <0.001 |
| WT/WT | 120 | parent[e] | 109 | 11 | | | | | | |
| *ERA8/ERA8* | 120 | parent[e] | 10 | 110 | | | | | | |
| WT/*ERA8* | 8 | $BC_2F_1$ | 4 | 4 | | | | | | |
| $F_3$ Expected[d] | 7361 | | | | 4409 | 2885 | 3655 | | | |
| 49d AR \| 5uM ABA \| 5d imb. | | | | | | | | | | |
| **X5.2b** | 583 | $BC_3F_3$ | 340 | 243 | 30.8 | 23.5 | 0.84 | <0.001 | <0.001 | 0.36 |
| WT/WT | 59 | parent[e] | 59 | 0 | | | | | | |
| *ERA8/ERA8* | 52 | parent[e] | 9 | 43 | | | | | | |
| WT/*ERA8* | 8 | $BC_2F_1$ | 4 | 4 | | | | | | |
| $F_3$ Expected[d] | 583 | | | | 402 | 282 | 329 | | | |
| 56d AR \| 10uM ABA \| 4d imb. | | | | | | | | | | |
| **X5.3**[f] | 1453 | $BC_3F_3$ | 344 | 1109 | 427.5 | 102.8 | 272.2 | <0.001 | <0.001 | <0.001 |
| WT/WT | 54 | parent[e] | 39 | 15 | | | | | | |
| *ERA8/ERA8* | 60 | parent[e] | 9 | 51 | | | | | | |
| WT/*ERA8* | 8 | $BC_2F_1$ | 4 | 4 | | | | | | |
| $F_3$ Expected[d] | 1453 | | | | 738 | 530 | 657 | | | |

[a] Number of seeds tested for germination, after-ripened for 5 weeks past physiological maturity; df = 1

[b] Generation of seeds tested

[c] Number of seeds that had germinated (Germ.) and not germinated (Not Germ.) after 5 days of imbibition on 2µM (+/-) ABA.

[d] Number of seeds expected to **germinate** after 5 days of imbibition for each single gene segregation ratio.

[e] Zak WT (+/+) and Zak *ERA8* (-/-) parental lines used to generate cross 5; Grown at the same time as the $F_1$ plants.

[d] Note that X5.3 was not used in any analyses since it appears to not have a 1:2:1 segregation.

**Table S6** Summary of[SAM1] the Louise/Zak*ERA8* RIL population GBS linkage groups

| Chrm | No. Groups [a] | Total Number of Markers | Map Distance (cM) |
|---|---|---|---|
| 1A | 1 | 231 | 534 |
| 1B | 3 | 147 | 334, 41, 88 |
| 1D | 2 | 57 | 73, 193 |
| 2A | 2 | 98 | 46, 272 |
| 2B | 1 | 49 | 270 |
| 2D | 1 | 57 | 483 |
| 3A | 4 | 180 | 296, 89, 33, 118 |
| 3B | 1 | 64 | 241 |
| 3D | 2 | 46 | 97, 153 |
| 4A | 3 | 224 | 279, 134, 30 |
| 4B | 2 | 130 | 294, 55 |
| 4D | 1 | 11 | 122 |
| 5A | 2 | 62 | 543, 42 |
| 5B | 3 | 84 | 138, 117, 38 |
| 5D | 2 | 27 | 239, 113 |
| 6A | 2 | 169 | 323, 168 |
| 6B | 2 | 39 | 167, 72 |
| 6D | 3 | 34 | 62, 167, 23 |
| 7A | 2 | 238 | 458, 272 |
| 7B | 2 | 206 | 227, 40 |
| 7D | 3 | 61 | 268, 78, 116 |
| unknown | 1 | 20 | 105 |
| total | 45 | 2,234 | - |

[a] Linkage groups were assigned to chromosomes based on the majority of markers aligned to one chromosome on the RefSeqv1.0 wheat genome (IWGSC 2018).

**Table S7** Genes in the *ERA8* interval on chromosome 4A between SNP_20 and SNP_29 markers

| Gene Model Name | Annotation | Start Position | Orientation | EMS SNP | Primer |
|---|---|---|---|---|---|
| *TraesCS4A01G299700* | High affinity cationic amino acid transporter 1, Uncharacterized protein | 597,908,536 | + | C → T | SNP_6 |
| *TraesCS4A01G311100* | Poly(A) RNA polymerase GLD2-A, Uncharacterized protein | 603,446,405 | | C → T | SNP_19 |
| *TraesCS4A01G311800* | Protein UPSTREAM OF FLC | 603,496,846 | - | - | |
| *TraesCS4A01G311900* | Antimicrobial peptide MBP-1 related (LEM1) | 603,504,474 | + | - | |
| *TraesCS4A01G312000* | TRAF-like superfamily protein | 603,507,001 | + | - | |
| *TraesCS4A01G312100* | Hydroxyproline-rich glycoprotein family protein | 603,510,936 | - | - | |
| *TraesCS4A01G312200* | GSK1 transcription factor 1 | 603,530,872 | - | | |
| | | 603,532,130 | | C → T | SNP_20 |
| *TraesCS4A01G312300* | Calcium-dependent lipid-binding domain protein | 603,533,711 | - | - | |
| *TraesCS4A01G312400* | AT hook motif DNA-binding family protein | 603,539,435 | - | - | |
| *TraesCS4A01G312500* | LOB domain-containing protein, putative | 603,634,586 | - | - | |
| *TraesCS4A01G312600* | Peptidyl-prolyl cis-trans isomerase | 603,637,101 | + | - | |
| *TraesCS4A01G312700* | Aquaporin-like protein | 603,666,065 | + | - | |
| *TraesCS4A01G312800* | Disease resistance protein (NBS-LRR class) family | 603,674,385 | + | - | |
| *TraesCS4A01G312900* | Disease resistance protein (NBS-LRR class) family | 603,698,121 | + | - | |
| *TraesCS4A01G313000* | Retrovirus-related Pol polyprotein from transposon TNT 1-94 | 603,699,642 | - | - | |
| *TraesCS4A01G313100* | Leucine rich repeat | 603,726,600 | - | - | |
| *TraesCS4A01G313200* | YUCCA9 | 603,980,902 | - | - | |
| *TraesCS4A01G313300* | PM19-A1 | 604,055,487 | - | - | |
| *TraesCS4A01G313400* | PM19-A2 | 604,068,519 | + | - | |
| *TraesCS4A01G313500* | Myosin-J-Protein | 604,076,529 | + | - | |
| *TraesCS4A01G313600* | Ubiquintin Congujating Enzyme E2-23 | 604,101,314 | + | - | |
| *TraesCS4A01G313700* | ACC Oxidase -1 Like | 604,104,116 | - | - | |
| *TraesCS4A01G313800* | B3-domain-containing protein | 604,107,899 | + | - | |
| *TraesCS4A01G313900* | LRR receptor-Like Kinase Serine/threonine-protein Kinase | 604,209,020 | - | - | |
| - | pseudogene | 604,315,235 | | C → T | SNP_33 |
| *TraesCS4A01G314000* | ACC Oxidase -1 Like | 604,594,773 | - | - | |
| *TraesCS4A01G314100* | Anthocyanin 5-aromatic acyltransferase | 604,659,700 | + | - | |
| *TraesCS4A01G314200* | LRR receptor-like kinase | 604,672,359 | + | - | |
| *TraesCS4A01G314300* | Disease resistance protein RPM1 | 604,772,917 | + | - | |
| *TraesCS4A01G314400* | Disease resistance protein RPM1 | 604,849,387 | - | - | |
| *TraesCS4A01G314500* | ERF-1B-Like | 604,856,919 | + | - | |
| *TraesCS4A01G314600* | ERF-1B-Like | 604,940,912 | + | - | |
| *TraesCS4A01G314700* | BTB/POZ/MATH-domain protein | 605,018,735 | - | - | |
| *TraesCSU01G167000* [a] | TaMKK3-A | - | | G → A | SNP_17 |
| *TraesCS4A01G314800* | transcription factor, putative (Protein of unknown function, DUF547) | 605,024,348 | + | - | |
| *TraesCS4A01G314900* | Sugar transporter, putative | 605,087,172 | - | - | |
| *TraesCS4A01G315000* | Serine/threonine-protein phosphatase | 605,137,125 | - | - | |
| - | low confidence gene | 605,375,951 | | G → A | SNP_34 |
| *TraesCS4A01G315100* | Protein kinase | 605,559,326 | - | - | |
| *TraesCS4A01G315200* | OTU domain-containing protein | 605,640,606 | + | - | |

| | | | | | |
|---|---|---|---|---|---|
| *TraesCS4A01G315300* | Trihelix transcription factor | 605,644,924 | + | - | |
| *TraesCS4A01G315400* | Early nodulin-like protein | 605,650,522 | - | - | |
| *TraesCS4A01G315500* | 60 kDa chaperonin | 605,656,215 | + | - | |
| *TraesCS4A01G315600* | Mediator of RNA polymerase II transcription subunit 15a | 605,663,094 | - | - | |
| *TraesCS4A01G315700* | RNA polymerase sigma factor | 605,711,154 | + | - | |
| *TraesCS4A01G315800* | Mediator of RNA polymerase II transcription subunit 13 | 605,717,570 | + | - | |
| *TraesCS4A01G315900* | ABC transporter G family member | 605,744,452 | + | - | |
| *TraesCS4A01G316000* | Cytoplasmic polyadenylation element-binding protein 4 | 606,116,294 | + | - | |
| *TraesCS4A01G316100* | F-box family protein | 606,335,446 | + | - | |
| *TraesCS4A01G316200* | Heavy metal-associated protein | 606,345,914 | - | - | |
| *TraesCS4A01G316300* | La-related protein | 606,361,307 | + | - | |
| *TraesCS4A01G316400* | GDSL esterase/lipase | 606,370,282 | + | - | |
| *TraesCS4A01G316500* | GDSL esterase/lipase | 606,397,403 | + | - | |
| *TraesCS4A01G316600* | Protein DETOXIFICATION | 606,411,251 | + | - | |
| *TraesCS4A01G316700* | Avr9/Cf-9 rapidly elicited protein | 606,523,442 | + | - | |
| *TraesCS4A01G316800* | WD40 repeat-containing protein | 606,536,121 | - | - | |
| *TraesCS4A01G316900* | phosphotransferases/inositol or phosphatidylinositol kinase | 606,585,463 | + | - | |
| *TraesCS4A01G317000* | Bifunctional inhibitor/lipid-transfer protein/seed storage 2S albumin-like protein | 606,591,102 | + | - | |
| *TraesCS4A01G317100* | DNA translocase FtsK | 606,593,057 | - | - | |
| *TraesCS4A01G317200* | Harpin-induced protein 1 (Hin1), putative | 606,608,730 | + | - | |
| *TraesCS4A01G317300* | Galactose-6-phosphate isomerase subunit LacB | 606,643,500 | - | - | |
| *TraesCS4A01G317400* | Mannonate dehydratase | 606,647,800 | - | - | |
| *TraesCS4A01G317500* | Kinase family protein | 606,761,092 | + | - | |
| *TraesCS4A01G317600* | Kinase family protein | 606,796,989 | + | - | |
| *TraesCS4A01G317700* | Oxidoreductase/transition metal ion-binding protein | 606,811,790 | + | - | |
| *TraesCS4A01G317800* | Exportin-1 | 607,045,327 | + | - | |
| *TraesCS4A01G317900* | Cytokinin riboside 5'-monophosphate phosphoribohydrolase | 607,177,351 | - | - | |
| *TraesCS4A01G318000* | Plant invertase/pectin methylesterase inhibitor superfamily | 607,178,696 | + | - | |
| *TraesCS4A01G318100* | Cytokinin riboside 5'-monophosphate phosphoribohydrolase | 607,261,062 | - | - | |
| *TraesCS4A01G318200* | F-box only 46 | 607,270,018 | + | - | |
| *TraesCS4A01G318300* | Leucine-rich repeat receptor-like protein kinase family protein | 607,270,941 | - | - | |
| *TraesCS4A01G318400* | ATP-dependent zinc metalloprotease FtsH | 607,309,139 | + | - | |
| *TraesCS4A01G318500* | DNA topoisomerase | 607,374,419 | - | - | |
| *TraesCS4A01G318600* | EMBRYO SURROUNDING FACTOR 1-like protein 8 | 607,378,421 | + | - | |
| *TraesCS4A01G318700* | Pentatricopeptide repeat-containing protein At1g19720 | 607,417,280 | + | - | |
| *TraesCS4A01G318800* | carbohydrate esterase, putative (DUF303) | 607,427,390 | - | - | |
| *TraesCS4A01G318900* | NAD/NADP-dependent betaine aldehyde dehydrogenase | 607,432,144 | + | - | |
| *TraesCS4A01G319000* | PGR5-like protein 1A, chloroplastic | 607,633229 | + | - | |
| - | - | 607,886,990 | | - | Barc170 |
| *TraesCS4A01G319100* | Gibberellin 20 oxidase | 608,043,459 | | G → A | SNP_29 |
| *TraesCS4A01G325400* | Pentatricopeptide repeat-containing protein; Pyridine nucleotide-disulphide oxidorectuse | 613,268,437 | | G → A | SNP_30 |

[a] The current reference genome (RefSeq v1.0; IWGSC 2018) aligned *TaMKK3-A* to the unknown chromosome. The estimated location of *TaMKK3-A* on chromosome 4A is based on Shorinola et al. (2017).

**Table S8** ABA sensitivity for the Louise/Zak*ERA8* RIL population was tested across three environments

| Location | Year | n[a] | t rep [b] | ABA (μM) | AR (days) | Mean ± SD [c] | | | | | |
|---|---|---|---|---|---|---|---|---|---|---|---|
| | | | | | | Day 1 | Day 2 | Day 3 | Day 4 | Day 5 | GI |
| Greenhouse | 2013 | 225 | 3 | 5 | 49 | 48.9 ± 29 | 71.9 ± 25 | 76.0 ± 22 | 78.8 ± 21 | 81.2 ± 19 | 0.71 ± 0.2 |
| (E1) | | *ERA8* | 9 | 5 | 49 | 26.7 ± 21 | 68.9 ± 21 | 74.4 ± 18 | 78.1 ± 16 | 81.1 ± 16 | 0.66 ± 0.2 |
| | | *Louise* | 9 | 5 | 49 | 70.4 ± 12 | 81.9 ± 13 | 86.3 ± 12 | 88.9 ± 10 | 91.9 ± 8 | 0.84 ± 0.1 |
| Field | 2014 | 181 | 3 | 2 | 42 | 22.3 ± 19 | 52.5 ± 25 | 63.3 ± 25 | 69.8 ± 25 | 74.9 ± 25 | 0.56 ± 0.2 |
| (E2) | | *ERA8* | 3 | 2 | 42 | 0.0 ± 0 | 5.6 ± 5 | 13.3 ± 12 | 17.8 ± 13 | 20.0 ± 12 | 0.11 ± 0.1 |
| | | *Louise* | 3 | 2 | 42 | 24.3 ± 8 | 44.4 ± 13 | 69.8 ± 18 | 77.8 ± 22 | 87.0 ± 11 | 0.58 ± 0.2 |
| Field | 2015 | 190 | 3 | 2 | 48 | 23.1 ± 22 | 71.5 ± 27 | 78.2 ± 24 | 82.2 ± 23 | 84.8 ± 21 | 0.68 ± 0.2 |
| (E3) | | *ERA8* | 3 | 2 | 48 | 5.6 ± 2 | 37.8 ± 23 | 43.3 ± 30 | 50.0 ± 27 | 53.3 ± 30 | 0.38 ± 0.2 |
| | | *Louise* | 3 | 2 | 48 | 27.8 ± 4 | 81.1 ± 10 | 86.7 ± 6 | 87.8 ± 7 | 88.9 ± 5 | 0.75 ± 0.1 |

[a] number (n) of recombinant inbred lines (RIL) tested

[b] technical replicates per RIL or parental line

[c] Raw mean and standard deviation (SD) across the RIL population, Louise, or *ERA8*.

**Table S9** Significant QTL in the Louise/Zak*ERA8* RIL population conducted with both the GBS and EMS-induce SNP markers

| QTL Name | Marker | Chrm | Pos (cM) | Start [a] | End [a] | SNP Position | LOD | Trait [b] | Favorable Allele [c] |
|---|---|---|---|---|---|---|---|---|---|
| *QABA.wsu-4A.1* | A11335 | 4A | 271.8 | 83,469,184 | 83,469,281 | 85 | 3.97 | E3 D1 | C/**T** |
| - | A14187 | 4A | 321.6 | 195,745,016 | 195,745,115 | 31 | 4.01 | E2 D2 | C/**T** |
| - | A13781 | 4A | 320.8 | 195,745,016 | 195,745,085 | 31 | 3.66 | E2 D2 | C/**T** |
| *QABA.wsu-4A.5* | A12726 | 4A | 295.9 | 216,478,520 | 216,478,451 | 47 | 4.8 | E2 D1 | G/**A** |
| *QABA.wsu-4A.5* | A10010 | 4A | 301 | 316,676,795 | 316,676,894 | 34 | 5.16 | E2 D1 | G/**A** |
| *QABA.wsu-4A.5* | A9539 | 4A | 301.8 | 316,676,795 | 316,676,864 | 34 | 5.02 | E2 D1 | G/**A** |
| *QABA.wsu-4A.5* | A25632 | 4A | 298.3 | 509,248,865 | 509,248,964 | 88 | 5.12 | E2 D1 | A/**G** |
| *QABA.wsu-4A.5* | A14030 | 4A | 299.7 | 526,310,792 | 526,310,693 | 49 | 4.19 | E2 D1 | C/**T** |
| *QABA.wsu-4A.5* | A25258 | 4A | 299.1 | 534,315,662 | 534,315,563 | 16 | 4.73 | E2 D1 | T/**C** |
| *QABA.wsu-4A.5* | A15895 | 4A | 297.3 | 534,325,792 | 534,325,693 | 47, 87 | 4.98 | E2 D1 | G, C / **C, T** |
| *QABA.wsu-4A.2* | SNP_5 | 4A | 325.5 | - | - | - | 4.45 | E2 D2 | G/**A** |
| *QABA.wsu-4A.2* | SNP_9 | 4A | 326 | 533,446,137 | - | - | 4.23 | E2 D2 | G/**A** |
| *QABA.wsu-4A.2* | SNP_4 | 4A | 326.2 | 532,074,134 | - | - | 4.32 | E2 D2 | C/**T** |
| *QABA.wsu-4A.2* | SNP_10 | 4A | 328.4 | 586,452,395 | - | - | 4.41 | E2 D2 | C/**T** |
| - | A18189 | 4A | 323.3 | 566,902,964 | 566,903,063 | 95 | 3.68 | E2 D2 | A/**G** |
| *QABA.wsu-4A.6* | A16514 | 4A | 294.7 | 573,801,680 | 573,801,581 | 52 | 4.31 | E2 D1 | C/**T** |
| *QABA.wsu-4A.6* | A16172 | 4A | 294 | 573,801,680 | 573,801,611 | 52 | 4.31 | E2 D1 | C/**T** |
| *QABA.wsu-4A.6* | A2917 | 4A | 292.5 | 577,057,065 | 577,056,996 | 49 | 4.67 | E2 D1 | G/**C** |
| *QABA.wsu-4A.8* | A29481 | 4A | 342.8 | 595,374,988 | 595,374,889 | 28 | 4.17 | E3 GI, D3 | T/**C** |
| *QABA.wsu-4A.8* | A29272 | 4A | 342 | 595,374,988 | 595,374,919 | 28 | 3.49 | E3 GI | T/**C** |
| *QABA.wsu-4A.8* | A22222 | 4A | 338.2 | 602,250,029 | 602,249,930 | 64 | 3.69 | E3 GI | G/**A** |
| *QABA.wsu-4A.4* | SNP_20 | 4A | 353.5 | 603,532,130 | - | - | 4.22 | E1 D1 | **G**/A |
| *QABA.wsu-4A.4* | TaMKK3-A | 4A | 356.2 | - | - | - | 5.35 | E1 D1 | **C**/A |
| *QABA.wsu-4A.4* | SNP_17 | 4A | 356.2 | - | - | - | 5.35 | E1 D1 | **G**/A |
| *QABA.wsu-4A.4* | SNP_29 | 4A | 358.7 | 608,044,262 | - | - | 4.44 | E1 D1 | **C**/T |
| *QABA.wsu-4A.10* | A190 | 4A | 439.1 | 623,366,977 | 623,366,908 | 17 | 4.07 | E3 D4-5, GI | **G**/T |
| *QABA.wsu-4A.7* | A28342 | 4A | 422.9 | 645,311,558 | 645,311,459 | 91 | 4.46 | E2 D2-5, GI | **A**/C |
| *QABA.wsu-4A.9* | A23553 | 4A | 141.1 | 658,854,423 | 658,854,492 | 55 | 3.96 | E2 D1 | **G**/A |
| *Qhd.wsu-4A* | A23913 | 4A | 50.93 | 658,854,423 | 658,854,522 | 55 | 4.87 | Heading | G/A |
| *Qhei.wsu-7B* | A2003 | 7B | 252.3 | 7,574,338 | 7,574,239 | 84 | 4.49 | Height | T/A |

[a] The GBS markers was aligned to the RefSeqv1.0 reference genome (IWGSC 2018) and the start and end nucleotide position is reported with the SNP position within the sequence.
[b] Significant QTL for the following traits are indicated by environment (E1, E2, or E3) followed by the percent germination after n days (D1-D5) of imbibition or germination index (GI). Significant QTL for heading date and height are also reported.
[c] Favorable alleles (underlined) decrease germination percent or index. Louise contributed the **first** and *ERA8* the **second** allele.

**Table S10** Unique genes differentially expressed between WT and *ERA8.* Highly differentially expressed genes are listed from the whole genome that were a) *ERA8* upregulate / WT downregulated (+ beta value) or b) *ERA8* downregulated / WT upregulated (- beta value). c) Differential expression of all unique genes located on chromosome 4A are listed with the EMS-induced SNP markers indicated in grey rows for reference.

| | Gene ID [a] | Annotation | Chrm | Start | End | beta [b] |
|---|---|---|---|---|---|---|
| **a)** | TraesCS2B01G524400 | - | 2B | 718,960,478 | 718,968,471 | **7.08** |
| | TraesCS3B01G322600 | - | 3B | 521,789,364 | 521,797,431 | **5.69** |
| | TraesCS2D01G000600 | Polycomb group protein VERNALIZATION 2 | 2D | 278,541 | 286,171 | **5.26** |
| | TraesCS6A01G338800 | DNA (Cytosine-5-)-methyltransferase | 6A | 572,133,054 | 572,138,876 | **5.20** |
| | TraesCS7D01G269200 | Reticulocyte-binding protein 2 a | 7D | 253,341,441 | 253,348,906 | **5.05** |
| **b)** | TraesCS7B01G229200 | - | 7B | 430,829,130 | 430,842,036 | **-5.20** |
| | TraesCS2D01G497100 | Protein UXT-like protein | 2D | 593,215,038 | 593,217,521 | **-5.24** |
| | TraesCS1A01G374800 | Zinc finger protein-like | 1A | 549,837,796 | 549,847,148 | **-5.39** |
| | TraesCS3B01G338100 | - | 3B | 544,780,837 | 544,793,535 | **-5.42** |
| | TraesCS7B01G484300 | DUF789 family protein | 7B | 741,573,444 | 741,579,514 | **-5.57** |
| | TraesCS2D01G500500 | - | 2D | 595,157,820 | 595,161,251 | **-5.58** |
| **c)** | TraesCS4A01G016000 | Transcription factor | 4A | 10,164,187 | 10,165,250 | -1.09 |
| | TraesCS4A01G021100 | F-box and associated interaction domains-containing protein | 4A | 14,340,239 | 14,344,018 | 1.47 |
| | TraesCS4A01G021900LC | Protein yippee-like | 4A | 17,974,706 | 17,977,539 | -0.87 |
| | TraesCS4A01G022000LC | Retrotransposon protein, putative, unclassified | 4A | 17,978,744 | 17,981,991 | **-1.80** |
| | TraesCS4A01G029300LC | Protein FAR1-RELATED SEQUENCE 5 | 4A | 25,211,087 | 25,215,287 | **4.17** |
| | TraesCS4A01G041700 | - | 4A | 35,181,518 | 35,183,648 | **4.29** |
| | TraesCS4A01G050500 | SNP_1 | 4A | 41,115,524 | - | - |
| | TraesCS4A01G052700LC | Sodium/hydrogen exchanger | 4A | 46,171,955 | 46,199,813 | 0.94 |
| | TraesCS4A01G052800LC | Sentrin-specific protease 1 | 4A | 46,173,178 | 46,174,094 | **2.92** |
| | TraesCS4A01G079100 | - | 4A | 80,840,552 | 80,844,726 | **4.10** |
| | TraesCS4A01G087400 | SNP_2 | 4A | 91,795,916 | - | - |
| | TraesCS4A01G092100 | Heat-shock protein, putative | 4A | 98,785,386 | 98,787,839 | 0.80 |
| | TraesCS4A01G097400 | Tryptophan synthase alpha chain | 4A | 108,369,445 | 108,371,437 | -1.43 |
| | TraesCS4A01G099900 | Histone H2B | 4A | 112,767,092 | 112,767,812 | -0.51 |
| | TraesCS4A01G103900 | SNP_13 | 4A | 117,337,422 | - | - |
| | TraesCS4A01G121600 | SNP_14 | 4A | 150,302,500 | - | - |
| | TraesCS4A01G126600 | 8-amino-7-oxononanoate synthase | 4A | 163,345,723 | 163,347,867 | -1.01 |
| | TraesCS4A01G130600 | NAC domain protein, | 4A | 173,630,224 | 173,632,115 | -0.69 |
| | TraesCS4A01G131600 | SNP_7 | 4A | 175,858,520 | - | - |
| | TraesCS4A01G131700 | SNP_3 | 4A | 176,543,353 | - | - |
| | TraesCS4A01G167000LC | Mitochondrial transcription termination factor family protein | 4A | 207,976,194 | 207,976,616 | 1.32 |
| | TraesCS4A01G179300LC | 2'-phosphotransferase | 4A | 230,997,232 | 231,003,186 | -0.67 |
| | TraesCS4A01G172100 | SNP_8 | 4A | 436,828,695 | - | - |
| | TraesCS4A01G248100LC | S-adenosyl-L-methionine-dependent methyltransferases superfamily protein | 4A | 340,489,977 | 340,490,856 | **2.02** |
| | TraesCS4A01G160600 | Phosphatidate phosphatase, Lipin | 4A | 345,835,861 | 345,844,921 | -1.17 |
| | TraesCS4A01G316400LC | Serine/threonine-protein kinase | 4A | 465,545,872 | 465,547,035 | 0.81 |
| | TraesCS4A01G323300LC | Cysteine desulfurase, putative, expressed | 4A | 474,726,464 | 474,739,302 | -0.47 |
| | TraesCS4A01G194800 | Non-specific serine/threonine protein kinase | 4A | 476,977,837 | 476,980,583 | -0.69 |
| | TraesCS4A01G202600 | Carboxypeptidase | 4A | 492,530,884 | 492,534,660 | -0.70 |

| | | | | | |
|---|---|---|---|---|---|
| TraesCS4A01G340900LC | MLO-like protein | 4A | 496,577,022 | 496,578,105 | **2.55** |
| TraesCS4A01G220100 | DNA polymerase | 4A | 522,907,385 | 522,921,948 | -0.92 |
| TraesCS4A01G221500 | Sphingoid base hydroxylase 2 | 4A | 527,112,529 | 527,113,641 | **2.65** |
| TraesCS4A01G224300 | SNP_4 | 4A | 532,074,134 | - | - |
| TraesCS4A01G225500 | SNP_9 | 4A | 533,446,137 | - | - |
| TraesCS4A01G232700, or TraesCS4A01G232800 | SNP_15 | 4A | 542,033,771 | - | - |
| TraesCS4A01G243200 | RING/U-box superfamily protein | 4A | 553,310,509 | 553,314,175 | 0.79 |
| TraesCS4A01G252800 | Cathepsin B-like cysteine protease | 4A | 565,040,316 | 565,043,816 | -1.32 |
| TraesCS4A01G411100LC | Transposon protein, putative, mutator sub-class | 4A | 566,101,663 | 566,104,738 | **3.76** |
| TraesCS4A01G258100 | Lariat debranching enzyme | 4A | 570,981,287 | 570,985,759 | 0.39 |
| TraesCS4A01G259300 | Anthocyanin 5-aromatic acyltransferase | 4A | 572,302,274 | 572,303,653 | 0.65 |
| TraesCS4A01G264600 | UDP-glucose 6-dehydrogenase | 4A | 576,930,165 | 576,932,752 | -0.60 |
| TraesCS4A01G278800 | SNP_10 | 4A | 586,452,395 | - | - |
| TraesCS4A01G444900LC | Peptide transporter | 4A | 591,496,571 | 591,499,205 | -0.97 |
| TraesCS4A01G288000 | Argonaute protein | 4A | 593,194,838 | 593,200,481 | 0.99 |
| TraesCS4A01G290300 | Ankyrin repeat family protein | 4A | 594,180,282 | 594,186,698 | 1.12 |
| TraesCS4A01G299700 | SNP_6 | 4A | 597,908,536 | - | - |
| TraesCS4A01G311100 | SNP_19 | 4A | 603,446,405 | - | - |
| TraesCS4A01G312200 | SNP_20 GSK1 transcription factor 1 | 4A | 603,532,130 | - | - |
| TraesCSU01G167000 | SNP_17 MKK3 | unk | - | - | - |
| - | barc170 | 4A | 607,886,990 | - | - |
| TraesCS4A01G319100 | SNP_29 GA 20-ox | 4A | 608,044,262 | - | - |
| TraesCS4A01G485400LC | BED zinc finger,hAT family dimerization domain | 4A | 612,510,456 | 612,511,014 | -1.26 |
| TraesCS4A01G485500LC | BED zinc finger,hAT family dimerization domain | 4A | 612,511,090 | 612,513,756 | **-2.60** |
| TraesCS4A01G325400 | SNP_30 | 4A | 613,286,437 | - | - |
| TraesCS4A01G331900 | Cinnamoyl-CoA reductase 4 | 4A | 616,309,725 | 616,313,994 | 0.44 |
| TraesCS4A01G345800 | Threonine synthase 1, chloroplastic | 4A | 625,029,910 | 625,030,335 | 0.86 |
| TraesCS4A01G359200 | - | 4A | 632,150,159 | 632,155,404 | 0.80 |
| TraesCS4A01G370500 | ABC transporter G family member | 4A | 642,209,939 | 642,213,850 | -0.94 |
| TraesCS4A01G386400 | Auxin repressed/dormancy associated protein | 4A | 663,986,105 | 663,987,373 | 0.59 |
| TraesCS4A01G593300LC | 3'(2'),5'-bisphosphate nucleotidase 1 | 4A | 674,553,838 | 674,556,604 | 0.69 |
| TraesCS4A01G401700 | Cysteine synthase | 4A | 675,600,394 | 675,603,117 | 0.53 |
| TraesCS4A01G599200LC | Zinc finger MYM-type protein 4 | 4A | 677,402,029 | 677,403,093 | 1.26 |
| TraesCS4A01G404800 | arabinogalactan protein 5 | 4A | 678,340,392 | 678,341,039 | -0.78 |
| TraesCS4A01G421900 | Histone H3 | 4A | 692,358,749 | 692,359,852 | -0.43 |
| TraesCS4A01G454800 | Glutathione S-transferase | 4A | 718,848,586 | 718,850,423 | 0.52 |
| TraesCS4A01G473900 | F-box protein | 4A | 733,670,552 | 733,672,307 | 1.01 |
| TraesCS4A01G485900 | Beta-fructofuranosidase 1 | 4A | 739,310,199 | 739,314,035 | **-1.50** |
| TraesCS4A01G486600 | disease resistance family protein / LRR family protein | 4A | 739,627,540 | 739,630,396 | **-1.64** |

[a] The genes model with LC are low confidence gene model and those without LC are High confidence gene models.

[b] Beta value is analogous to fold-change in slueth. A fold change of an absolute value of 1.5 or higher is considered differentially expressed and indicated in **bold**. Highly differentially expressed genes" correspond to those with beta > |5|. A positive fold-change means there was more expression in *ERA8* compared to WT. A negative fold-change means there was more expression in WT compared to *ERA8*. Highly differentially expressed genes" correspond to those with beta > |5|
